# Protein complex prediction for large protein protein interaction networks with the Core&Peel Method

**DOI:** 10.1101/053876

**Authors:** Marco Pellegrini, Miriam Baglioni, Filippo Geraci

## Abstract

**Motivations.:** Biological networks play an increasingly important role in the exploration of functional modularity and cellular organization at a systemic level. Quite often the first tools used to analyze these networks are *clustering algorithms*. We concentrate here on the specific task of predicting protein complexes (PC) in large protein-protein
interaction networks (PPIN). Currently, many state-of-the-art algorithms work well for networks of small or moderate size. However, their performance on much larger networks, which are becoming increasingly common in modern proteome-wise studies, needs to be re-assessed. Our aim is to push forward the state-of the-art in PPIN clustering providing an algorithmic solution with polynomial running time that attains experimentally demonstrable good output quality and speed on challenging large real networks.

**Results.:** We present a new fast algorithm for clustering large sparse networks: *Core&Peel*, which runs essentially in time and storage *O*(*a*(*G*)*m*+*n*) for a network *G* of *n* nodes and *m* arcs, where *a*(*G*) is the arboricity of *G* (which is roughly proportional to the maximum average degree of any induced subgraph in *G*). We evaluated *Core&Peel* on five PPI networks of large size and one of medium size from both yeast and homo sapiens, comparing its performance against those of ten state-of-the-art methods. We demonstrate that *Core&Peel* consistently outperforms the ten competitors in its ability to identify known protein complexes and in the functional coherence of its predictions. Our method is remarkably robust, being quite insensible to the injection of random interactions. *Core&Peel* is also empirically efficient attaining the second best running time over large networks among the tested algorithms.

**Availability:** http://bioalgo.iit.cnr.it (via web interface)

## 1 Introduction

Due to recent advances in high-throughput proteomic techniques, such as yeast two-hybrid system (Y2H) and Tandem Affinity Purification coupled with Mass Spectrometry (TAP-MS), it is now possible to compile large maps of protein interactions, which are usually denoted as *protein-protein interaction networks* (PPIN). However, extracting useful knowledge from such networks is not straightforward. Therefore sophisticated PPI network analysis algorithms have been devised in the last decade for several goals such as: the prediction of protein-complexes ([1]), the prediction of higher level functional modules ([2, 3, 4]), the prediction of unknown interactions ([5, 6]), the prediction of single protein functions ([7]), the elucidation of the molecular basis of diseases ([8]), and the discovery of drug-disease associations ([9]), to name just a few. In this paper we concentrate on the issue of predicting protein-complexes (PC) in PPI networks. An incomplete list of complex prediction algorithms in chronological order is: MCODE [10], RNSC [11], Cfinder [12], MCL [13], COACH [14], CMC [15], HACO [3], CORE [16], CFA [17], SPICi [18], MCL-CAw [19], ClusterONE [20], Prorank [21], the Weak ties method [22], Overlapping Cluster Generator (OCG) [23], PLW [24], PPSampler2 [25], and Prorank+ [26]. Further references to existing methods can be found in recent surveys by [27], [28], [29], [1], [30], and [31].

The graph representing a PPIN can also be augmented so to include additional biological knowledge, annotations and constraints. The conservation of protein complexes across species as an additional constraint is studied in [32]. Jung et al. [33] encode in PPIN the information on mutually exclusive interactions. Proteins in PPIN can also be marked with cellular localization annotations ([34]), and several types of quality scores. Though all these aspects are important, they are possible refinements applicable to the majority of the algorithms listed above, involving the modeling of additional knowledge in the PPIN framework (see [35]). In this paper we concentrate on the basic case of a PPIN modeled as an undirected and unweighted graph. The size of PPIN found in applications tend to grow over time because one can obtain with modern techniques from a single high-throughput experiment thousands of novel PPI, and also because one can collate groups of PPI from different experiments into a single larger network (ensemble PPIN) [36]. For example very large PPIN arise in multi-species PPI studies, ([37], [38]), in immunology studies ([39, 40]) and cancer data analysis ([41]). Large PPIN can be challenging for clustering algorithms as many of them have been designed and tested in the original publication with PPIN of small and medium size (with the possible exception of SPICi ([18]), that was designed intentionally for large PPIN). Greedy methods that optimize straightforward local conditions may be fast but speed may penalize quality. Thus, although more than a decade has passed since the first applications of clustering to PPIN, the issue of growing PPIN size poses new challenges and requires a fresh look at the problem.

We develop a new algorithm (*Core&Peel*) designed for clustering large PPIN and we apply it to the problem of predicting protein complexes in PPIN. The complexes we seek have just very basic properties, they should appear within the PPIN as ego-networks of high density and thus we can model them as maximal quasi-cliques. These features are not particularly new, but we show in section 5 that they are sufficient to characterize a large fraction of PCs in a sample of five large PPIN for two species (yeast and human). Computational efficiency is attained by a systematic exploitation of the concept of *core decomposition* of a graph, which for each vertex (protein) in a graph provides a tight upper bound to the size of the largest quasi-clique that includes that vertex. We use this upper bound to trim locally the subgraphs of interest in order to isolate the sought quasi-clique, and proceed then to the final *peeling* out of loosely connected vertices. Our approach has some superficial similarity with that of CMC ([15]) which applies the enumeration algorithm of [42] to produce, as an intermediate step, a listing of all maximal cliques in a graph.

We avoid this intermediate step that may cause an exponential running time in large PPIN and cannot be adapted easily to listing all maximal quasi-cliques, when density below 100% is sought. Our approach is both more direct (no intermediate listing of potentially exponential size is produced) and more flexible (as we can tune freely and naturally the density parameter).

CFinder ([12]) lists all k-cliques, for a user defined value of *k*, and then merges together k-cliques sharing a (k-1)-clique. CFinder might produce too many low density clusters if the user choosees k too small, or miss interesting complexes if k is too large. *Core&Peel* avoids both pitfalls since we have a more adaptive control over cluster overlaps. Our algorithm is empirically very fast: all instances in this paper run in less than 2 minutes on common hardware. The asymptotic analysis (see Additional_file_1 Section 8) indicates a running time very close to linear for sparse graphs. More precisely, with some additional mild sparsity assumptions, the algorithm runs in time *O*(*a*(*G*)*m* + *n*) for a graph *G* of *n* nodes and *m* arcs, where *a*(*G*) is the *arboricity* of *G* (which is roughly proportional to the maximum average degree of any induced subgraph in *G*). The output quality is assessed by comparative measures of the ability to predict known complexes and of the ability to produce biologically homogeneous clusters, against 10 state-of-the art methods. In both quality assessments *Core&Peel* leads or ties in most tests vs all other methods, often by a large margin (See Section 6). The robustness of our method is remarkably high, since practically no output variation is measured even when adding up to 25% random edges in the input graph. Finally, we show several high quality predicted clusters that involve a known complex with additional proteins, which correspond to biologically relevant mechanisms described in literature.

## 2 Paper Organization

In Section 3 we start by reviewing the issue of false positive/negative PPI in large PPIN with hindsight from the work in [5] indicating quasi-cliques as good models for protein complexes in our settings (Section 3.1). Next, in Section 3.2 we recall the basic graph-theoretic definitions of subgraph density, quasi-cliques, and core-decompositions, that are central to our algorithmic design. In Section 3.3 we introduce the notion of a *partial dense cover* as a formalization of our problem, showing its similarities with well known NP-hard problems of *minimum clique cover* and *maximum clique* [43]. In Section 3.4 we give a high level description of our proposed polynomial time heuristic. For ease of description it is split in four phases, though in optimized code some of the phases may be interleaved. The rationale behind certain design choices is explained in further detail in Section 3.5. The asymptotic analysis of the proposed algorithm can be found in Additional_file_1 (Section 8).

The experimental set up is described in Section 4, including the sources of raw data, the initial data cleaning (Section 4.1) and the quality score functions (Sections 4.2 and 4.3). Further data statistics and details of the comparative evaluations are in Section 5 and 6. In particular we report on the ability to capture known complexes in Section 6.1, to produce functionally coherent clusters (Section 6.2), on robustness in presence of random noise (Section 6.3), and on computation timings (Section 6.4).

In Section 6.5 we list ten interesting predictions in which a known complex interacts with an additional protein. These findings have an independent support in the literature. Finally in Section 7 we comment on the potential applications and extensions of the proposed method, as well as on its limitations.

## 3 Method

### 3.1 On false positive and false negative PPI in dense and large PPIN

The estimation of the number of erroneous PPI calls (false positive/false negative) in PPI networks is highly dependent on the technology and the experimental protocols used. Yu et al. [5] report an experiment on 56 proteins of Saccharomyces cerevisiae (yeast) for which PPI were detected using both error-prone high throughput technologies and more precise low throughput technologies. In 563 cases (pairs of proteins) for which the two methods differ, the vast majority (92.5%) were false negatives (FN), and just 7.5% false positive (FP). A similar ratio among FP/FN rates is reported in [36] for PPI obtained through Y2H and high confidence AP-MS techniques. While each technology has its own systematic biases, it is observed in [36] that such biases tend to compensate each other when data from several sources is used to compile ensemble PPIN. The implication is that, over time, as the evidence on reliable PPI accumulates, the number of undetected real PPI (FN) will steadily decrease, while the number of spurious PPI (FP) should increase quite slowly. In graph terms the subgraphs representing complexes in the PPI will become denser (i.e. closer to a clique), while the noisy interactions will still remain within a controllable level (assuming that only high quality interaction data is encoded in the PPI networks). Expanding on these finding Yu et al. [5] demonstrate that *quasi-cliques*(cliques with a few missing edges) are good predictors of the presence of a protein complex, provided the PPIN is large. Our own measurents on one medium size graphs (≈ 20*K* PPI) and four large graphs (≈ 130*K/*220*K* PPI) in Section 5 confirm this tendency of protein complex density increase in larger PPIN. Besides the increase in density, a second notable phenomenon, is that protein complexes often resemble ego-networks, that is, the protein complex is mostly contained in the 1-neighborhood of some protein (see Section 5).

### 3.2 Preliminaries

An early incarnation of the *Core&Peel* algorithm targeting communities in social graphs is described in [44]. In order to make this paper self-contained we are describing in this section a version of *Core&Peel* that includes all the modifications needed to target potentially overlapping protein complexes in PPI network. Let *G* = (*V*, *E* ⊆ *V* × *V*) be a simple (undirected) graph (no self-loops, no multiple edges). A subset *Q* ⊂ *V* induces a subgraph *H*_*Q*_ = (*Q,E*_*Q*_), where *E*_*Q*_ = {(*a*, *b)* ∈ *E*\*a* ∈ *Q* ∧ *b* ∈ *Q*}. For a graph *G* its *average degree* is:

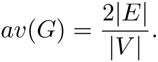

The *density* of a graph *D*(*G*) is the following ratio:

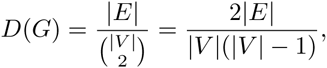

which gives the ratio of the number of edges in *G* to the maximum possible number of edges in a complete graph with the same number of nodes. We restrict ourselves to *local density definitions*, such as the two listed above, that are those for which the density of a subgraph induced by a subset *Q* ⊆ *V* is a function depending only on *Q* and on the induced edges set *E*_*Q*_. A nice survey of concepts and algorithms related to local density of subgraphs is in [45]. Cliques are subgraphs of density 1, and finding a maximum induced clique in a graph *G* is an NP-complete problem [46]. Several relaxations of the notion of clique have been proposed (see [47] for a survey), most of which also lead to NP-complete decision problems. Given a parameter γ ∈ [0..1], a γ-quasi clique is a graph *G = (V, E)* such that:

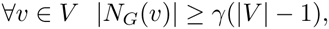

where *N*_*G*_(*v*) = {*u* ∈ *V* |(*v*, *u*) ∈ *E*} is the set of immediate neighbors of *v* in *G*. Note that a γ-quasi clique has density *D*(*G*) ≥ γ. In general, however, for a dense graph with density *D(G)* we cannot infer a bound on the value of γ for which there exists a quasi-clique in G (except for the value *D*(*G*) = 1 that implies γ = 1, and those cases covered by Turán’s theorem ([48])). If we impose that the number of vertices in a subgraph is exactly *k*, then the average degree and the density depend only on the number of edges, and thus they attain their maximum values for the same subgraphs. Without this constraint, finding the subgraph of maximum average degree or the subgraph of maximum density are quite different problems: the former admits a polynomial time solution, the latter is NP-complete. In this paper we aim at detecting dense-subgraphs with a lower bound on the size of each sub-graph and on its density, thus still an NP-complete problem. A *k-core* of a graph *G* is a maximal connected subgraph of *G* in which all vertices have degree at least k. A vertex u has *core number* k if it belongs to a k-core but not to any (k+1)-core. A *core decomposition* of a graph is the partition of the vertices of a graph induced by their core numbers ([49]).

### 3.3 Partial dense cover of a graph

In this section we formalize our problem as that of computing a *partial dense cover* of a graph. We aim at collecting efficiently only high quality candidate dense sets that cover the dense regions of the input graph. A Partial Dense Cover *PDC*(*G*, *r*, *δ,q*) is defined as the range of the function ƒ: *V* → 2^*V*^ that associates to any vertex *v* ∈ *V* a subset of *V* with these properties:

a. if ƒ(*v*) ≠ ∅ then *v* ∈ ƒ(*v*), (the set ƒ(*v*) contains the seed *v* or it is empty).
b. ƒ(*v*) ⊆ *N* ^*r*^(*v*) ⋃ {*v*}, (the set ƒ(*v*) is a subset of the *r*-neighborhood of *v*, i.e. all its vertices are at distance at most *r* from *v*. (In this study, we set *r* = 1 throughout)
c. ƒ(*v*) is the largest set having size at least *q*, density at least *δ*, satisfying (a) and (b), or otherwise it is the empty set.

Note that there may be more than one set ƒ(*v*) that, for a given *v*, satisfies (a), (b) and (c). If this is the case, we pick arbitrarily one such set as the value of ƒ(*v*). Since the *PDC*(*δ,q*)^[1]^ is the range of the function ƒ, by definition, it contains no duplicate sets, though its elements can be highly overlapping. One way to imagine this structure is as a relaxation of a *minimum clique cover* of a graph that is the problem of determining the minimum value *k* such that the vertices of a graph can be partitioned into *k* cliques. We relax this problem by (1) relaxing the disjointness condition (we allow sets to overlap) (2) allowing also a covering with graphs of density smaller than 1.0 (cliques correspond to density value *δ* = 1.0). Computing a clique cover of minimum size *k* is a well known NP-complete problem [50], and it is hard to approximate [51]. Even in this weaker form it remains NP-complete, by an easy reduction to the *maximum clique problem*. The cover we seek is *partial* since we do not insist that every vertex must be included in some set. We exclude sets that are too small (below a size threshold *q*) or too sparse (below a density threshold *δ*). The size parameter *q* and density parameter *δ* ensure that we can focus the computational effort towards those part of the graph that are more interesting (i.e. of large size and high density) with the goal of attaining computational efficiency while collecting high quality dense candidate sets. Note that for *δ* = 1.0 the *PDC*(1.0, *q*) is a subset of the set of all maximal cliques. While the set of all maximal cliques can be much larger than |*V* |, actually a worst case exponential number [43, 52], the *PDC*(*δ,q*) has always at most |*V* | elements (and in practical cases quite fewer than that).

### 3.4 Algorithm *Core&Peel* in highlight

As noted above, computing a partial dense cover of a graph is a NP-complete problem. In this section we describe an efficient heuristic algorithm which is based on combining in a novel way several algorithmic ideas and procedures already presented separately in the literature. For each step we give intuitive arguments about its role and an intuitive reason for its contribution to solving the problem efficiently and effectively. We first give a concise description of the four main phases of the *Core&Peel* algorithm. Subsequently we describe each phase in more detail.

#### Algorithm Overview

##### Phase I

Initially we compute the *Core Decomposition* of *G* (denoted with *CD*(*G*)) using the linear time algorithm in [53], giving us the core number *C*(*v*) for each node *v* ∈ *V*. Moreover we compute for each vertex *v* in *G* the *Core Count* of *v*, denoted with *CC*(*v*), defined as the number of neighbors of *v* having core number at least as large as *C(v)*. Next, we sort the vertices of *v* in decreasing lexicographic order of their core values *C*(*v*) and core count value *CC*(*v*).

##### Phase II

In Phase II we consider each node *v* in turn, in the order given by Phase I. For each *v* we construct the set *N*_*C*(*v*)_(*v*) of neighbors of *v* in *G* having core number greater than or equal to *C(v)*. We apply some filters based on simple node/edge counts in order to decide whether *v* should be processed in Phase III. If |*N*_*C*(*v*)_(*v*)| < *q* we do not process this node any more, being too small a set to start with. Otherwise we apply one of the following filters. We compute the density *δ*(*v*) of the induced subgraph *G*[*N*_*C*(*v*)_(*v*)]. If this density is too small (i.e. *δ*(*v*) ≤ *δ*_*low*_) for a threshold *δ*_*low*_, which we specify later, we do not process this node any more (filter (f=0)). In the second filter (f=1) we check if there are at least *q* nodes with degree at least(*q* - 1)*δ*. The third filter (f=2) is a combination of the previous two filters. Nodes that pass the chosen filter are processed in Phase III.

##### Phase III

In this phase we take *v* and the induced subgraph *G*[*N*_*C*(*v*)_(*v*)] and we apply a variant of the peeling procedure described in [54] that iteratively removes nodes of minimum degree in the graph. The peeling procedure stops (and reports failure) when the number of nodes drops below the threshold *q*. The peeling procedure stops (and reports success) when the density of the resulting subgraph is above or equal to the user defined threshold *δ*. The set of nodes returned by the successful peeling procedure is added to the output cover set.

##### Phase IV

Here we eliminate duplicates and sets completely enclosed in other sets, among those passing the Phase III. We also test the Jaccard coefficient of similarity between pairs of predicted complexes, removing one of the two predictions if they are too similar according to a user-defined threshold.

## 3.5 Algorithm description: Details

Many of our choices rely in part on provable properties of the core number and of the peeling procedure shown in [54], and in part on the hypothesis that the peeling procedure will converge to the same dense subgraph for both notions of density, when the initial superset of nodes is sufficiently close to the final subset. However the connections between these properties, the approximation to a partial dense cover computed by the algorithm, and the properties of validated protein complexes in a PPIN network can be only conjectured. The final justification of individual choices is mainly based on the good outcome of the experimental evaluation phase.

### Details on Phase I

The core decomposition of a graph *G* = (*V*, *E*) associates to any vertex *v* a number *C*(*v*) which is the largest number such that *v* has at least *C(v)* neighbors having core number at least *C*(*v*). Consider now a clique *K*_*x*_ of size *x*, for each node *v* ∈ *K*_*x*_ its core number is *x* - 1. If *K*_*x*_ is an induced subgraph of *G*, then its core number is at least *x* - 1, thus *C*(*v*) is an upper bound to the size of the largest induced clique incident to *v*. Consider a γ-quasi-clique *K*_(*x*,γ)_ of *x* nodes, for each node *v* in *K*(_*x*,γ_) its core number is at least γ(*x* - 1). If *K*_(*x*,γ)_ is an induced subgraph of *G*, then its core number can only be larger, thus *C*(*v*) is an upper bound to the size of the largest (in terms of average degree) quasi-clique incident to *v*. Thus if the upper bound provided by the core number is tight, examining the nodes in (decreasing) order of their core number allows us to detect first the largest cliques (or quasi-cliques), and subsequently the smaller ones.

In a clique *K*_*x*_ each node is a *leader* for the clique, meaning that it is at distance 1 to any other node in the clique. Thus the first node of *K*_*x*_ encountered in the order computed in Phase I is always a leader. In the case of quasi-cliques of radius 1 we have by definition the existence of at least one leader node. For an isolated quasi-clique the leader node will have the maximum possible core count value, thus by sorting (in the descending lexicographic order) on the core count value we force the leader node to be discovered first in the order (assuming all nodes in the quasi-clique have the same core number). For an induced quasi-clique the influence of other nodes may increase the value of the core count for any node, but, assuming that the relative order between the leader and the other nodes does not change, we still obtain the effect of encountering the leader before the other nodes of the quasi-clique.

The core number of a node *v* gives us an estimate of the largest (in terms of average degree) quasi-clique (or clique) incident to *v*, thus it provides a very powerful filter. We employ the very simple and very efficient algorithm in [53] that computes the core decomposition of a graph in time and storage *O*(|*V* | + |*E* |).

### Details on Phase II

In Phase II we aim at computing simple conditions and we decide whether node *v* should be processed in the next (more expensive) phase III. The first condition to test is |*N*_*C*(*v*)_(*v*)| < q, i.e. whether the number of nodes is below the user defined lower bound for the size (this is applied always). We apply then one of the following filter policies. We define the filter policy ƒ = 0, by checking a sufficient condition for the existence of a clique in a dense graph based on the classical results of Turán ([48]) that guarantees the existence of a clique (or a clique with a few edges missing) in graphs with sufficiently many edges. (approximately above *n*^2^/4 for a graph of *n* nodes). This corresponds to setting *δ*_*low*_ = 1/2, which indeed did perform well in our experiments with radius 1. We define the filter policy ƒ = 1, by checking the necessary condition for the existence of a *δ*-quasi clique of at least *q* nodes (this condition is that *G*[*N*_*C*(*v*)_(*v*)] must contain at least *q* nodes of degree at least (*q* - 1)*δ*). Finally, we define the filter policy ƒ = 2, that is the union of the previous two filters.

### Details on Phase III

The peeling procedure we use is similar to the one described in [54]. It consists in an iterative procedure that removes a node of minimum degree and all its incident edges, and iterates on the residual graph. In [54] the graph of highest average degree constructed in this process is returned as output. We modify this procedure by returning the first subgraph generated that satisfies the density and size constraints. It is shown in [54] that this procedure is (1/2)-approximate for the *maximum average degree*, i.e. it returns a subgraph whose average degree is within a factor 1/2 of that of the subgraph of highest average degree. Empirically, we rely on the intuition that the input to the peeling procedure produced after Phase II is a superset of the target dense subgraph and that it is sufficiently tight and dense so that the peeling procedure converges quickly and the target dense subgraph is isolated effectively. We also use a novel heuristic to solve cases of ties within the peeling algorithm in [54]. When two or more vertices are of minimum degree the original peeling procedure picks one arbitrarily. In our variant we compute the sum of degrees of the adjacent nodes *S*(*v*) = Σ_*w*∈*N*(*v*)_ |*N*(*w*)| and we select the vertex among those of minimum degree minimizing *S*(.). This secondary selection criterion is inspired by observations in [55], where the objective is to select an independent set by iteratively removing small degree nodes, which is a dual of the problem of detecting cliques.

### Details on Phase IV

In order to eliminate duplicate sets, we collect all the sets passing phase III, we split them in equal length classes and we represent them as lists of node identifiers in sorted order. Next we do a lexicographic order of each class, thus lists that are equal to each other end up as neighbors in the final sorted order and they can be easily detected and removed. In order to further exploit the sparsity of the output of phase III, we represent the collection of sets {Γ_*i*_} produced in phase III, with duplicates removed, as a graph whose nodes are the sets and elements of {Γ_*i*_}. The edges represent the inclusion relation. In this graph the number of 2-paths joining nodes Γ_*i*_ and Γ_*j*_ is exactly |Γ_*i*_ ∩ Γ_*j*_|. If |Γ_*i*_ ∩ Γ_*j*_| = |Γ_*j*_|, we know Γ_*j*_ ⊂ Γ_*i*_ and we can remove Γ*j*. We can count efficiently such number of 2-paths by doing a Breadth First Search at depth 2 starting from each set-node in the bipartite graph in increasing order of size, and by removing each starting node after its use. This operation allows us to compute if a set is a subset of another set, and also the Jaccard coefficient of similarity of any two non-disjoint sets.

## 4 Experimental set up

### 4.1 Used Data and preprocessing

We used the following freely accessible data sets to test our method.

#### Protein Protein interaction networks

Biogrid ([56]): we downloaded both Biogrid homo sapiens^[2]^ and Biogrid yeast^[3]^. String ([38]): we downloaded the general String file^[4]^ and then we extracted the two subsets of interest: the homo sapiens one (related to the 9606 NCBI taxonomy id) and the yeast one (related to the 4932 NCBI taxonomy id). DIP ([57]): we downloaded the yeast db^[5]^.

#### Protein databases

From the NCBI web site we downloaded the two files for homo sapiens^[6]^ and yeast^[7]^, the Uniprot db (uniprot_sprot.dat on 26/03/2013), and the Ensembl mapping for the associations of *ensemblproteinid* with *entrez id* for homo sapiens.

#### Protein Complexes

We downloaded CYC2008 ([58]) and CORUM ([59]) data on 26/03/2013.

#### Gene Ontology (GO)

We downloaded the files for homo sapiens^[8]^ on 10/09/2014, and for yeast^[9]^ on 10/09/2014

#### Preprocessing

Files from different sources of PPI are heterogeneous in many aspects. DIP exploits the *Uniprot accession id* (or other db entries as aliases) to represent the proteins involved in the interaction, Biogrid exploits the *NCBI entrez id*, and String uses *Ensembl proteins id* for homo sapiens and *gene locus* or *Uniprot accession* for yeast. The first operation was to represent in a uniform way the proteins for both the PPI files and the gold standard files. We decided to represent each protein with their associated NCBI entrez-id. In the process we removed possible duplications, and proteins for which the mapping was not possible. For the String data we also removed PPI with a quality score below 700. For the GO file, we identified and separated the three principal categories of the gene ontology, which are Cellular Component (CC), Biological Process (BP), and Molecular Functions (MF). Following the methodology in [20], these files are filtered to remove the annotation with IEA, ND and NAS evidence codes (corresponding to the “Inferred from electronic annotation”, “No biological data available” and “Non-traceable author statement”, respectively). Each protein associated to an annotated function is then mapped to its NCBI entrez id. Eventual repetitions of proteins for an annotation have been removed.

### 4.2 Evaluation measures for protein complex prediction

In order to better capture the nuisances of matching predicted clusters with actual complexes, we use four scalar measures (one from [28], and three from [60]) and we sum them to form a single scalar *Aggregated Score* (AS). Each of the four measures differs form the others in some key aspects: some use a step-function, while other use cluster-size as weights. All four, however, aim at balancing precision and recall effects. A similar aggregation of indices has been used in [20], although we use a different pool of indices.

#### F-measure

From [28] we adopted the following f-measure computation to estimate the degree of matching between the found cluster and the gold standard complex. Let *P* be the collection of discovered clusters and let *B* be the collection of the gold standard complexes. For a pair of sets *p* ∈ *P* and *b* ∈ *B*, the *precision-recall product score* is defined as *PR*(*p,b*) = 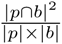. Only the clusters and complexes that pass a *PR*(*p*, *b*) threshold *ω* (step function) are then used to compute precision and recall measures. Namely we define the matching sets: *N*_*p*_ = |{*p* |*p* ∈ *P*, ∃*b* ∈ *B*, *PR*(*p*, *b*) ≥ *ω*}|, and *N*_*b*_ = |{*b* |*b* ∈ *B*, ∃*p* ∈ *P*, *PR*(*p*, *b*) ≥ *ω*}|. Afterwards: *Pecision* = 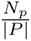, *Recall* = 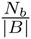, and the *F-measure* is the harmonic mean of precision and recall. In line with [28] and other authors we use *ω* = 0.2. Experiments in [61] indicate that the relative ranking of methods is robust against variations of the value of *ω*.

From [60] we adopted three measures to evaluate the overlap between complexes and predicted clusters: the *Jaccard measure*, the *precision-recall measure* and the *semantic similarity measure*.

#### Jaccard measure

Let the sets *P* and *B* be as above, for a pair of sets *p* ∈ *P* and *b* ∈ *B*, their Jaccard coefficient is *Jac*(*p,b*) = 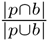. For each cluster *p* it is defined *Jac*(*p*) = *max*_*b*∈*B*_*Jac*(*p*, *b*), and for each complex *b* it is defined *Jac*(*b*) = *max*_*p*∈*P*_*Jac*(*p*, *b*). Next, we compute the weighted average Jaccard measures using, respectively, the cluster and complex sizes: *Jaccard*(*P*) = 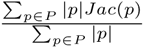, and *Jaccard*(*B*) = 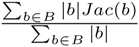. Finally, the *Jaccard measure* is the harmonic mean of*Jaccard*(*P*) and *Jaccard*(*B*).

#### Precision Recall Product

This measure is computed using exactly the same work flow as *Jaccard*, except that we replace the Jaccard coefficient with the precision-recall product score used also in [28].

#### Semantic similarity Measure

Let the sets *P* and *B* be as above, for a protein *x*, we define *P*(*x*) as the set of predicted clusters that contain *x*: *P*(*x*) = {*p* ∈ *P* |*x* ∈ *p*}, and *B*(*x*) as the set of golden complexes that contain *x*: *B*(*x*) = {*b* ∈ *B* |*x* ∈ *b*}. Denote with *I*(.) the indicator function of a set that is 0 for the empty set and 1 for any other set. Let *Bin*(.) denote the set of unordered pairs of distinct elements of a set. The semantic similarity of *p* in *B* is: *Den*(*p,B*) = 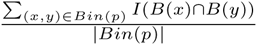. Analogously the semantic similarity of *b* in *P* is: *Den*(*b,P*) =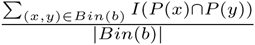. Next, we compute the weighted average semantic similarity weighted respectively by cluster and complex size: *Density*(*P*) = 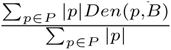, and *Density*(*B*) = 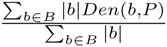. Finally, the *Semantic Similarity Measure* is computed as the harmonic mean of *Density*(*P*) and *Density*(*B*).

#### Handling of small protein complexes

The presence or absence of small protein complexes in the golden standard and in the outcome of the algorithms complicates the evaluation, thus in Additional_file_1 Section 4 we describe a fair method for placing all algorithms on a level field with respect to this issue.

### 4.3 Evaluation measure for Gene ontology coherence

For a predicted cluster *p* ∈ *P* we compute a **q-value** score trying to assess its biological coherence and relevance. Let *G* be a collection of gene ontology annotations, and *g* one GO class. Let *M* be the set of all proteins. For a predicted cluster *p*, we compute the *hypergeometric* p-value *H*(*M,p,g*) of the association of *p* to *g*, when *g* ∩ *p* ≠ ∅:

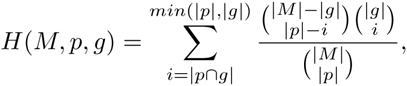

which represents the probability that a subset of *M* of size *|p|* chosen uniformly at random has with *g* an intersection of size larger than or equal to |*p* ∩ *g*|. As, in general, *p* will have an hypergeometric score for each gene ontology class it intersects, following [20] and [62], we associate to each *p* the intersecting gene ontology class of lower p-value. In order to correct for multiple comparisons we correct the vector of p-values using the q-value method of [63] which is a regularized version of the Benjamini Hochberg FDR estimation method. The q-values for the vector of p-values are computed via the R package provided at http://genomine.org/qvalue/.

## 5 Experiments

### Basic direct measures

Basic measures on the PPINs and protein complexes data sets are reported in Table 1 and in Table 2, respectively. When we map the known curated complexes onto the PPI-networks we obtain 5 different data sets in which the number and density of the embedded complexes is specific to the involved PPIN (see Table 3). The resulting embedded complexes have variable density. We report in Table 3 the 90% and the 50% density percentiles. One of the assumptions we have used in our algorithm is that for each embedded complex there is one vertex that is linked to (almost) all the other nodes in the embedded complex (egocentricity). This is an important property that measures on the actual data support (see Table 4). In Table 5 we report on the degree of overlap among complexes by counting the number of proteins belonging to one, two, three or more than three complexes. This is an important feature of the prediction problem since algorithms need to handle properly overlapping clusters. Human complexes have higher overlap rates than yeast complexes. In Additional_file_1 (section 6) we report the distributions of basic measures relative to the graph (degree, core number, clustering coefficients), and to the embedded PC (size, average degree, density).

**Table 1.** Columns give: PPI name, Species (Sp.)(hs=homo sapiens, y=yeast), reference, number of proteins |*V*|, number of interactions|*E*|, average degree 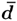, and whether a quality filter (Fil.) has been applied.

| PPI | Sp. | Ref. | $\vert V\vert $ | $\vert E\vert $ | $d$ | Fil. |
| --- | --- | --- | --- | --- | --- | --- |
| DIP | y | [57] | 4,637 | 21,107 | 9.1 | No |
| Biogrid | y | [56] | 6,686 | 220,499 | 65.9 | No |
| String | y | [38] | 5,590 | 133,082 | 47.6 | Yes |
| Biogrid | hs | [56] | 18,170 | 137,775 | 15.1 | No |
| String | hs | [38] | 12,717 | 193,105 | 30.3 | Yes |

**Table 2.** Columns give: name of the data set, Species (Sp.)(hs=homo sapiens, y=yeast), reference, total number of complexes, number of complexes of size 3 or larger, number of complexes of size up to 2, total number of proteins covered by the complexes.

| Name | Sp. | Ref. | # complexes | # compl. size $\geq 3$ | # compl. size $\leq 2$ | # proteins |
| --- | --- | --- | --- | --- | --- | --- |
| CYC2008 | y | [58] | 408 | 236 | 172 | 1627 |
| CORUM | hs | [59] | 1750 | 1257 | 493 | 2506 |

**Table 3.** Columns give: Name of the PPI and complex data set, number of complexes of size ≥ 3, min size, max size, average size, number of complexes with density *δ* greater than 0.9 and 0.5.

| Name | # CX | min | max | mean | $\delta > 0.9$ | $\delta > 0.5$ |
| --- | --- | --- | --- | --- | --- | --- |
| DIP-CYC2008 | 226 | 3 | 40 | 6.02 | 60 (25%) | 131 (55%) |
| BioGrid-CYC2008 | 236 | 3 | 81 | 6.67 | 173 (73%) | 223 (94%) |
| String-CYC2008 | 236 | 3 | 81 | 6.67 | 220 (93%) | 235 (99%) |
| BioGrid-CORUM | 1257 | 3 | 143 | 6.12 | 516 (41%) | 943 (75%) |
| String-CORUM | 1188 | 3 | 133 | 6.07 | 621 (52%) | 981 (82%) |

**Table 4.** Columns give: Name of the PPI and complex data set, number of complexes of size ≥ 3, number of complexes with at least one center at distance 1 (*r* = 1) for a fraction of at least 0.9 of its size and at least 0.5 of its size. Similar data for a center at distance 2, (*r* = 2).

| Name | # CX | $r_1 > 0.9$ | $r_1 > 0.5$ | $r_2 > 0.9$ | $r_2 > 0.5$ |
| --- | --- | --- | --- | --- | --- |
| DIP-CYC2008 | 226 | 131 (55%) | 197 (83%) | 163 | 212 |
| BioGrid-CYC2008 | 236 | 216 (91%) | 234 (99%) | 234 | 236 |
| String-CYC2008 | 236 | 235 (99%) | 236 (100%) | 236 | 236 |
| BioGrid-CORUM | 1257 | 891 (70%) | 1162 (92%) | 1176 | 1246 |
| String-CORUM | 1188 | 923 (77%) | 1139 (95%) | 1085 | 1188 |

**Table 5.**
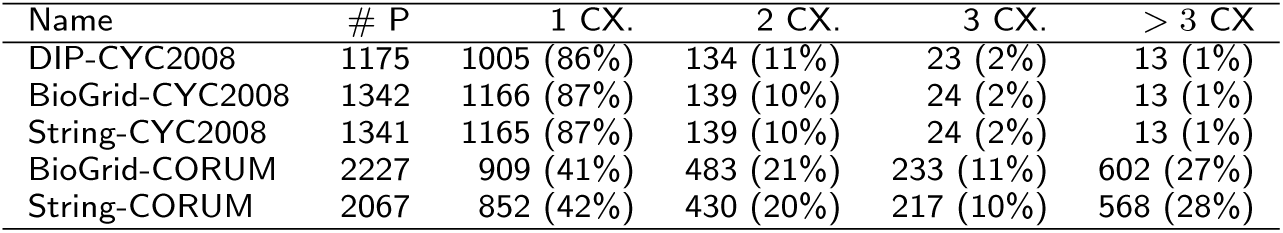
Columns give: Name of the PPI and complex data set. Number of proteins covered by some complex. Number of protein covered by one, two, three or more then three complexes.

| Name | # P | 1 CX. | 2 CX. | 3 CX. | $> 3$ CX |
| --- | --- | --- | --- | --- | --- |
| DIP-CYC2008 | 1175 | 1005 (86%) | 134 (11%) | 23 (2%) | 13 (1%) |
| BioGrid-CYC2008 | 1342 | 1166 (87%) | 139 (10%) | 24 (2%) | 13 (1%) |
| String-CYC2008 | 1341 | 1165 (87%) | 139 (10%) | 24 (2%) | 13 (1%) |
| BioGrid-CORUM | 2227 | 909 (41%) | 483 (21%) | 233 (11%) | 602 (27%) |
| String-CORUM | 2067 | 852 (42%) | 430 (20%) | 217 (10%) | 568 (28%) |

### Quality testing

We report the comparative evaluation of our algorithm vs several other algorithms, among those considered state-of-the-art. We used for these experiments an Intel core i7 processor (4 cores) at 2.6GHz, with 16Gb RAM memory, and with Mac OS X 10.8.5.

We have selected 10 algorithms^[10]^ among those in literature by applying these criteria: (a) we selected algorithms that appeared in several surveys and comparative evaluations, and well cited in the literature; (b) we included both old classical algorithms and more recent ones; (c) we have included algorithms using definitions of density similar to the one we adopt; (d) we included algorithms with available implementation in the public domain or obtainable from the authors upon request; (e) we preferred implementations based on widely available (i.e. non-proprietary) platforms; (f) we avoided algorithms that make use of additional biological annotations (e.g. gene expression data); (g) we preferred methods with a clear and unique underlying algorithm (e.g. “ensemble” methods are not included); (h) we preferred methods that aim at”protein complex detection” vs. those that aim at”functional module discovery”, since the evaluation methodologies for these two classes are quite different, although many methods could be construed as dual-use. Each method has its own pool of parameters to be set. For the quality score shown in section 4.2 we have considered for each method an extensive range of input parameter values (see File Additional_file_1, section 2 and 3) and we selected for each quality measure used in the Aggregated Score the best result obtained. Note that each best value for the four base quality measures may be obtained with slightly different values of the control parameters. Missing measures indicate that, for a specific algorithm and data set, the computation would not complete within a reasonable amount of time (without any sign of progress) or it generated fatal runtime errors.

## 6 Comparative evaluation

### 6.1 Performance of Protein Complex Prediction

Figures 1, 2, 3, 4 and 5 report the F-measure, the Semantic Similarity, the J-measure, the PR-measure and the Aggregated Score (as defined in Section 4.2) for three data sets relative to yeast PPIN (DIP, Biogrid and String). Out of 15 measurements, *Core&Peel* has the best value in 12 cases, CMC in 2 cases, and ClusterOne in 1 case. The Aggregated Score, which balances strong and weak points of the four basic measures, indicates that *Core&Peel*, CMC and ClusterOne have about the same performance for the medium-size PPI newtwork DIP. But for Biogrid data and even more for String data *Core&Peel* takes the lead, even with a wide margin. Figures 6, 7, 8, 9 and 10 report the F-measure, the Semantic similarity, the J-measure, the PR-measure and the aggregated score for three data sets relative to homo sapiens PPI (Biogrid and String). During the evaluation of the predicted clusters for Biogrid data we realized that the Biogrid PPI network had one node of very high degree corresponding to the Ubiquitin (UBC) protein. This fact has a straightforward biological explanation. Since UBC is involved in the degradation process of other proteins, UBC is linked to many other proteins at a certain time in their life-cycle. Given this special role of UBC, when protein degradation is not the main focus of the intended investigation, it may be convenient to consider also the same PPI network with the UBC node and its incident edges removed^[11]^ (labelled BG-hs-UBC). We tested also the other PPI network used in our study and this is the only case in which removing a node of maximum degree changes significantly the outcome of the prediction. Out of 15 measures, *Core&Peel* has the best value in all 15 cases. Good performance is obtained on some measures by CMC and Spici. It is interesting to notice how the algorithms perform differently on the BG-hs with and without UBC. On Biogrid data without UBC, *Core&Peel*, Spici and ClusterOne improve their AS value, while RNSC and COACH have a reduced AS value. The improvement in absence of UBC can be easily explained by the fact that UBC appears only in a few complexes of the golden standard, thus the evaluation phase is made more precise by its removal from the network and thus from the predicted clusters. The better results attained by RNSC and COACH on the graph with UBC may be a hint that, for these two approaches, the presence of UBC helps in homing in more quickly on the true complexes hidden in the graph.

**Figure 1.**
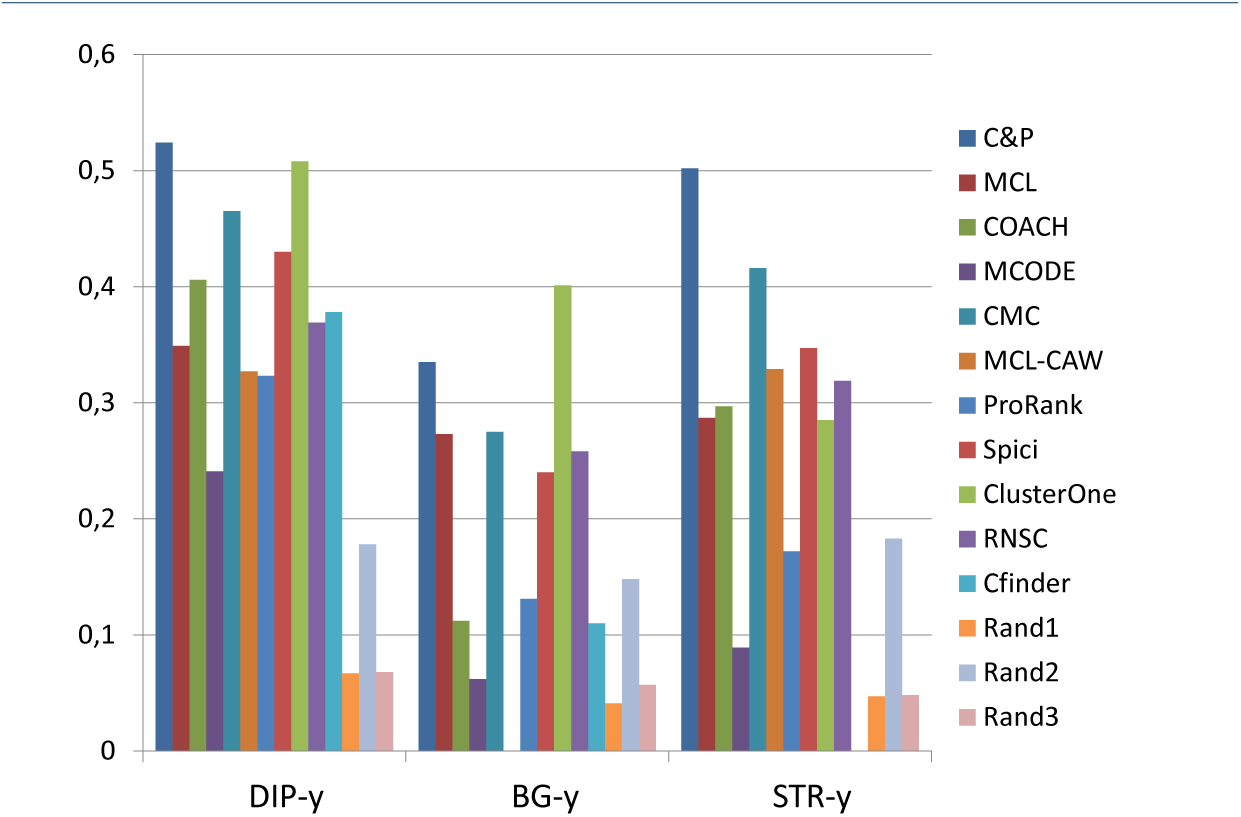
F-measure score for 11 algorithms and 3 random baselines on yeast data. Runs optimizing the f-measure for each algorithm.

**Figure 2.**
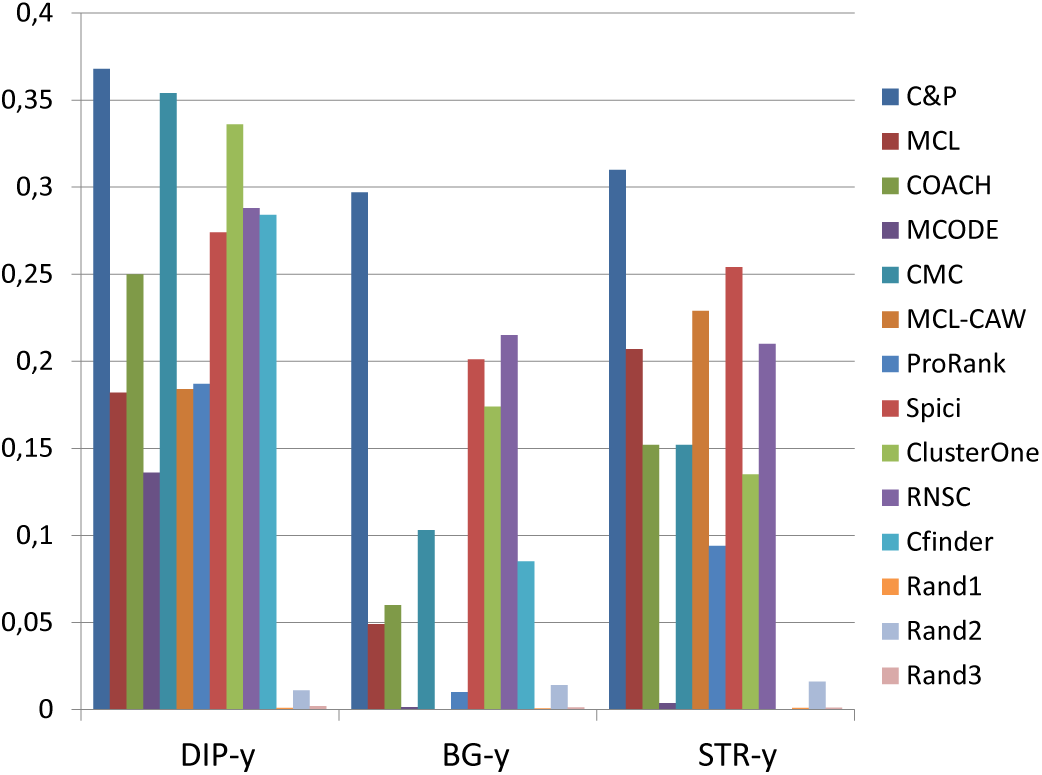
Semantic Similarity score for 11 algorithms and 3 random baselines on yeast data. Runs optimizing the ss-measure for each algorithm.

**Figure 3.**
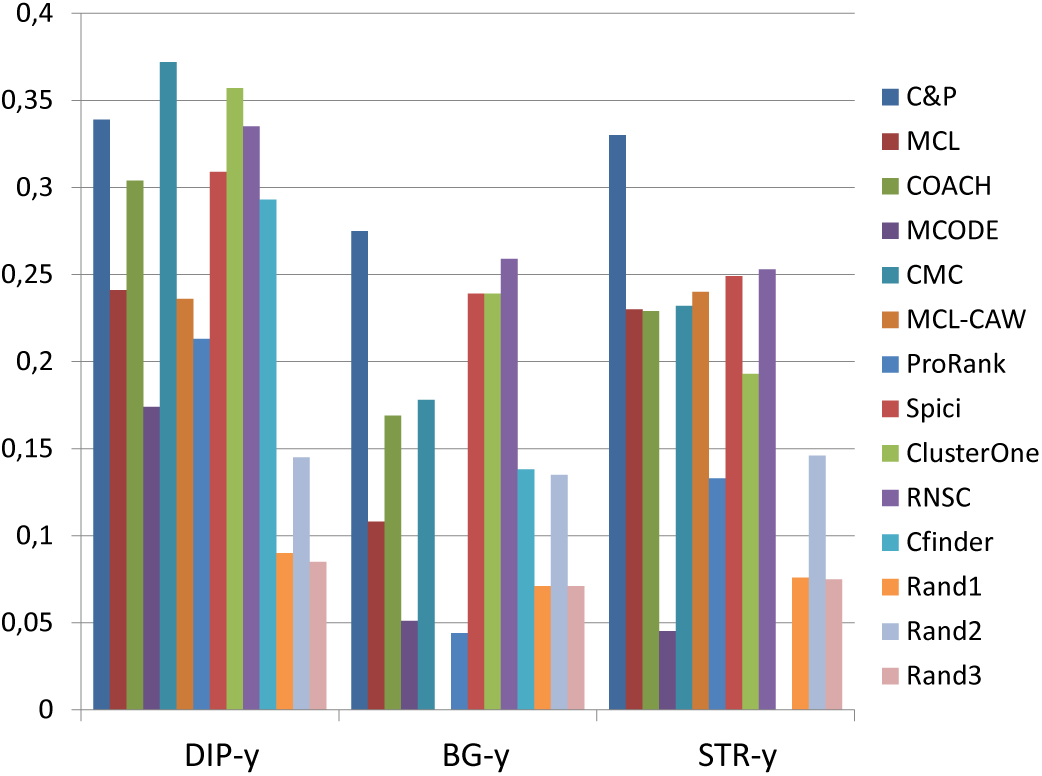
J-measure score for 11 algorithms and 3 random baselines on yeast data. Runs optimizing the J-measure for each algorithm.

**Figure 4.**
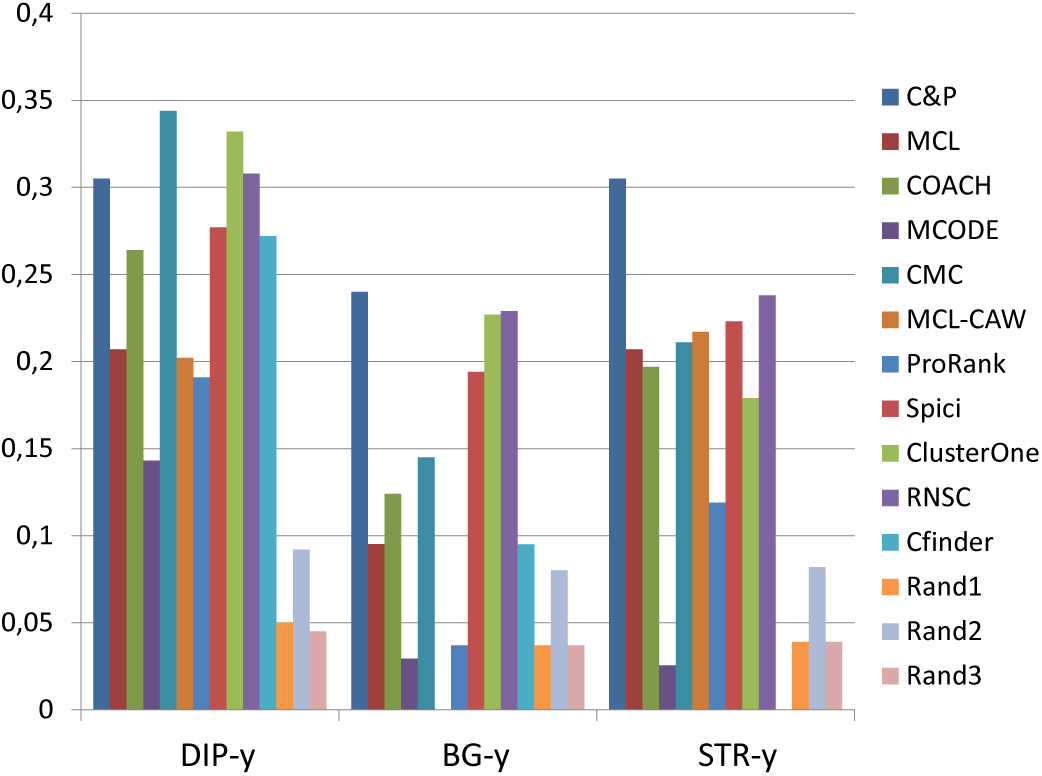
PR-measure score for 11 algorithms and 3 random baselines on yeast data. Runs optimizing the PR-measure for each algorithm.

**Figure 5.**
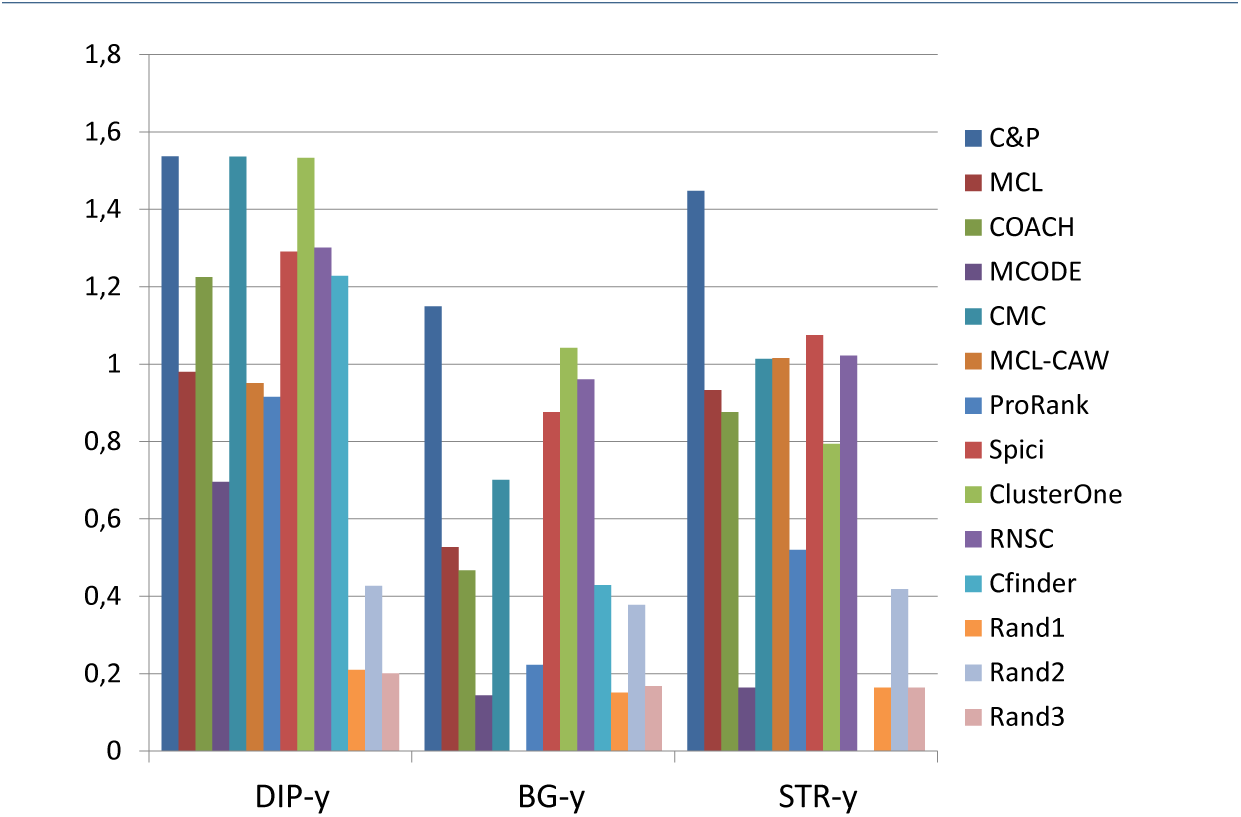
Aggregated score for 11 algorithms and 3 random baselines on yeast data.

**Figure 6.**
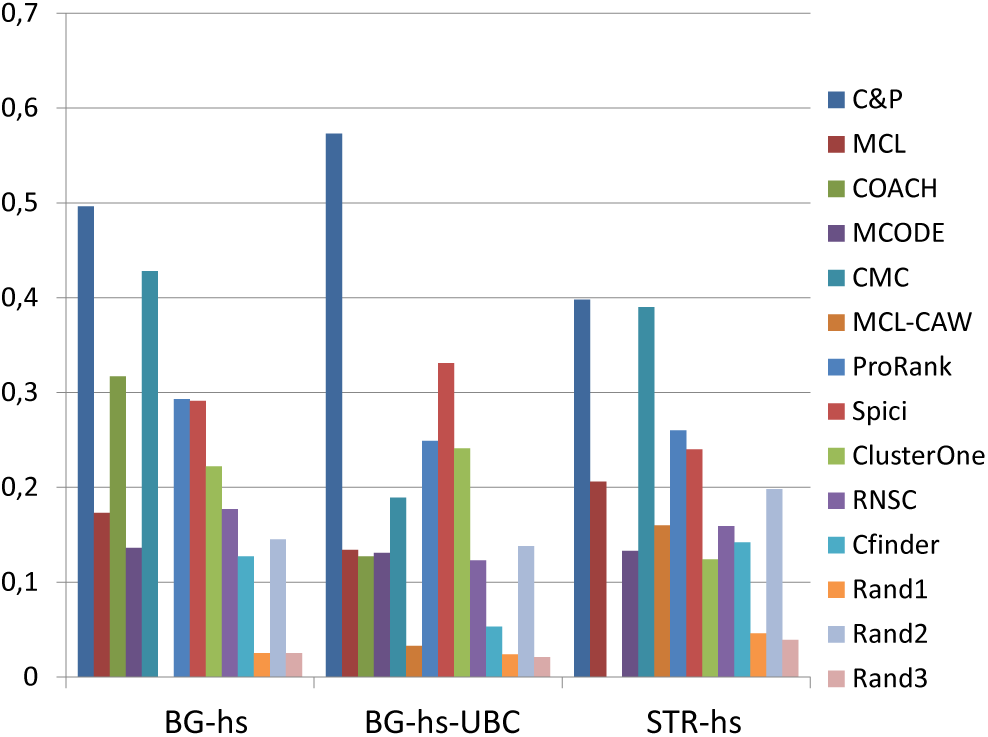
F-measure score for 11 algorithms and 3 random baselines on homo sapiens data. Runs optimizing the f-measure for each algorithm.

**Figure 7.**
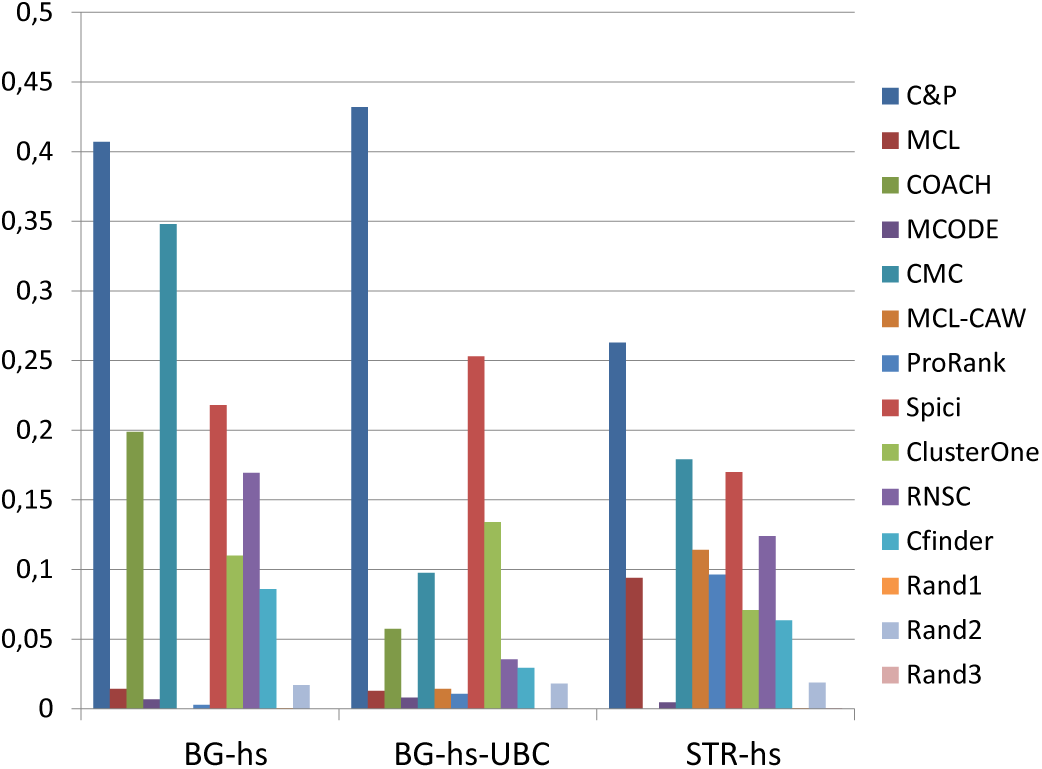
Semantic Similarity score for 11 algorithms and 3 random baselines on homo sapiens data. Runs optimizing the SS-measure for each algorithm.

**Figure 8.**
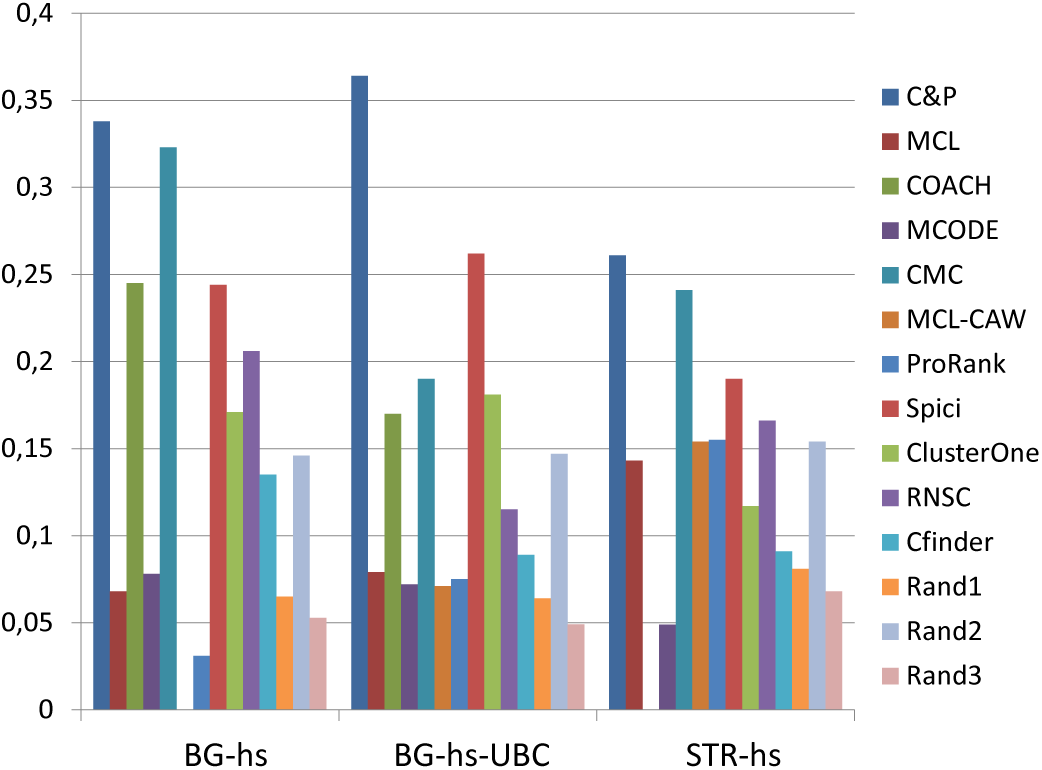
J-measure score for 11 algorithms and 3 random baselines on homo sapiens data. Runs optimizing the J-measure for each algorithm.

**Figure 9.**
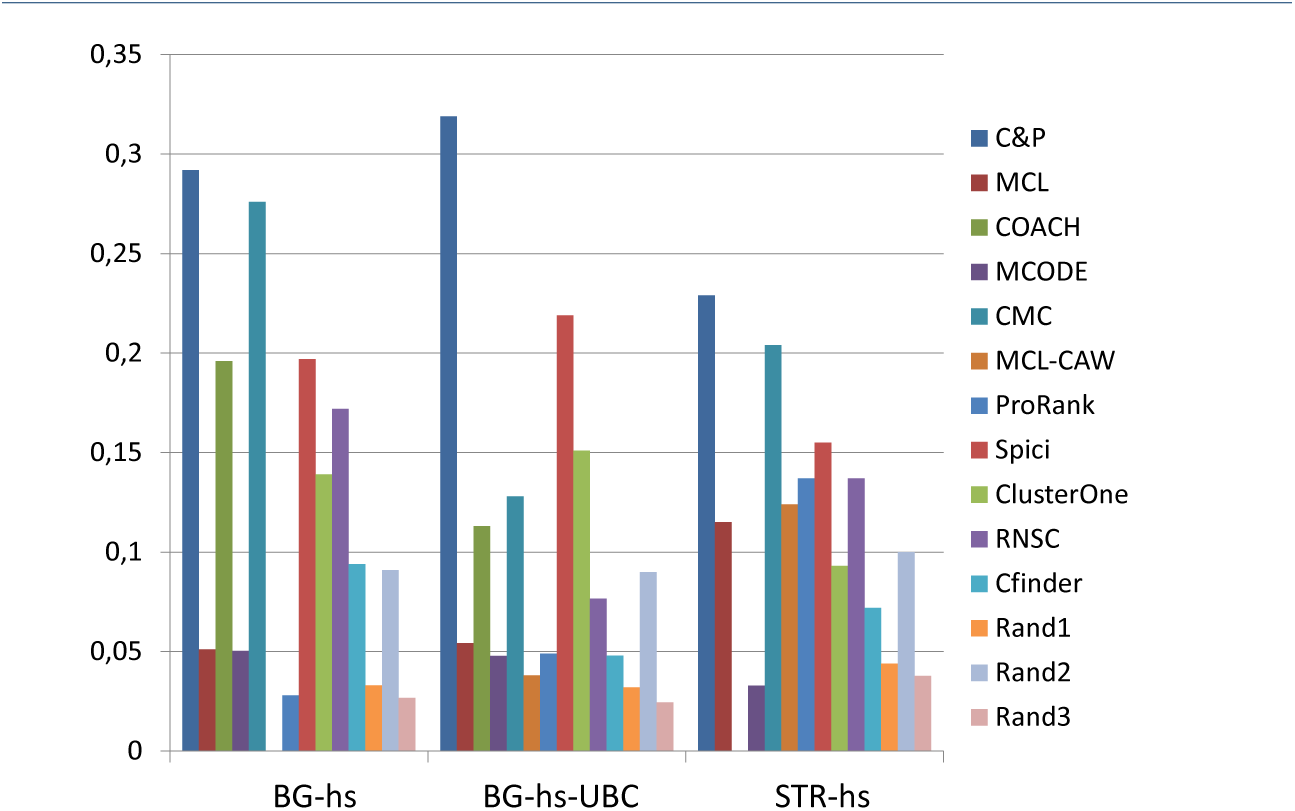
PR-measure score for 11 algorithms and 3 random baselines on homo sapiens data. Runs optimizing the PR-measure for each algorithm.

**Figure 10.**
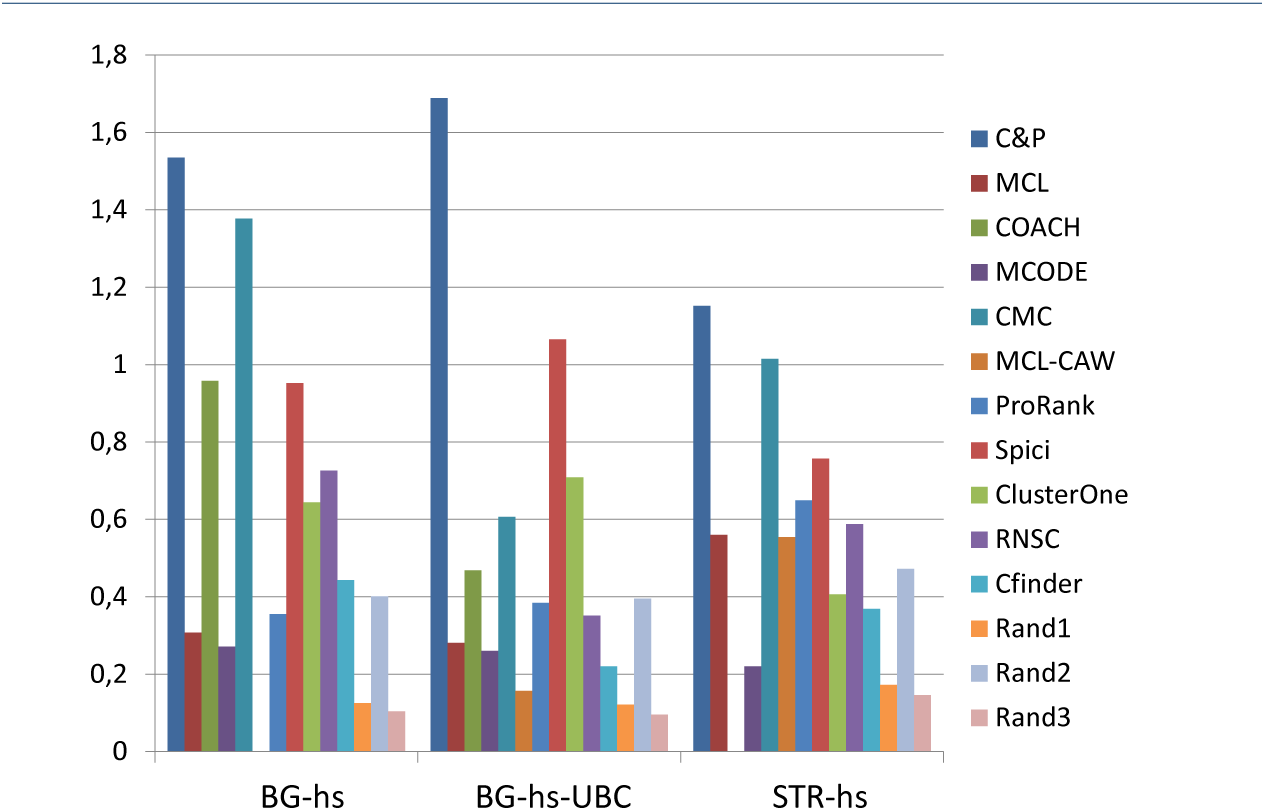
Aggregated score for 11 algorithms and 3 random baselines on homo sapiens data.

We include as a sanity check also three random predictions (Rand1, Rand2, and Rand3). The purpose of this check is to assess how well the measure we are using are able to discriminate the predictions on real data sets from those generated randomly by generators allowed to access some partial knowledge about the structure of the golden standard.

The method Rand1 is given the size distribution of the sets in the golden standard and produce a random collection of sets out of the vertices of the PPI with the same size distribution. The method Rand2 is as Rand1 except that the random sets are generated starting from the subset of all vertices in the PPI that belong to some complex in the golden standard.

The method Rand3 is obtained by taking the golden standard and applying to it a random permutation of the nodes of the PPI. Note that this approach besides preserving the size distribution preserves also the distribution of the size of the intersections of any number of sets of the golden standard.

In terms of performance, Rand1 behaves almost like Rand3, while Rand2 (having stronger hints) attains better results. The semantic similarity measure is the one that has better discrimination power vs all the three random test cases.

*Core&Peel* has better SS performance on all the 6 PPIN tested than the 10 competing methods. Semantic similarity is the only measure that explicitly places a premium in correctly identifying the proteins that simultaneously belong to multiple complexes, thus we can infer that *Core&Peel* successfully uncovers the overlapping structure of the the known protein complexes.

### 6.2 Coherence with Gene Ontology Annotation

The second index is the number of predicted clusters with an associated functional annotation (Biological Processes (BP) of Gene Onontology (GO)) below a given False Discovery Rate (FDR) threshold. Note that here we use a non-normalized measure (absolute count) since we want to favor algorithms with a rich high quality output. We are safeguarded against rewarding unduly methods that inflate their output since we operate each algorithm with the parameters that optimize the (balanced) F-measure. Moreover, even though none of the methods we use incorporates GO as part of its model, it is relatively safe to assume that, in most cases of interest, GO annotations are indeed available and may be used for a post-processing re-ranking or filtering of the predictions.

The biological function enrichment measure (using the BP annotation in GO) is shown in Figures 11, 12, 13, 14, 15, and 16. We used in abscissa the FDR thresholds ranging from 10^−2^ to 10^−7^ on the q-value.

**Figure 11.**
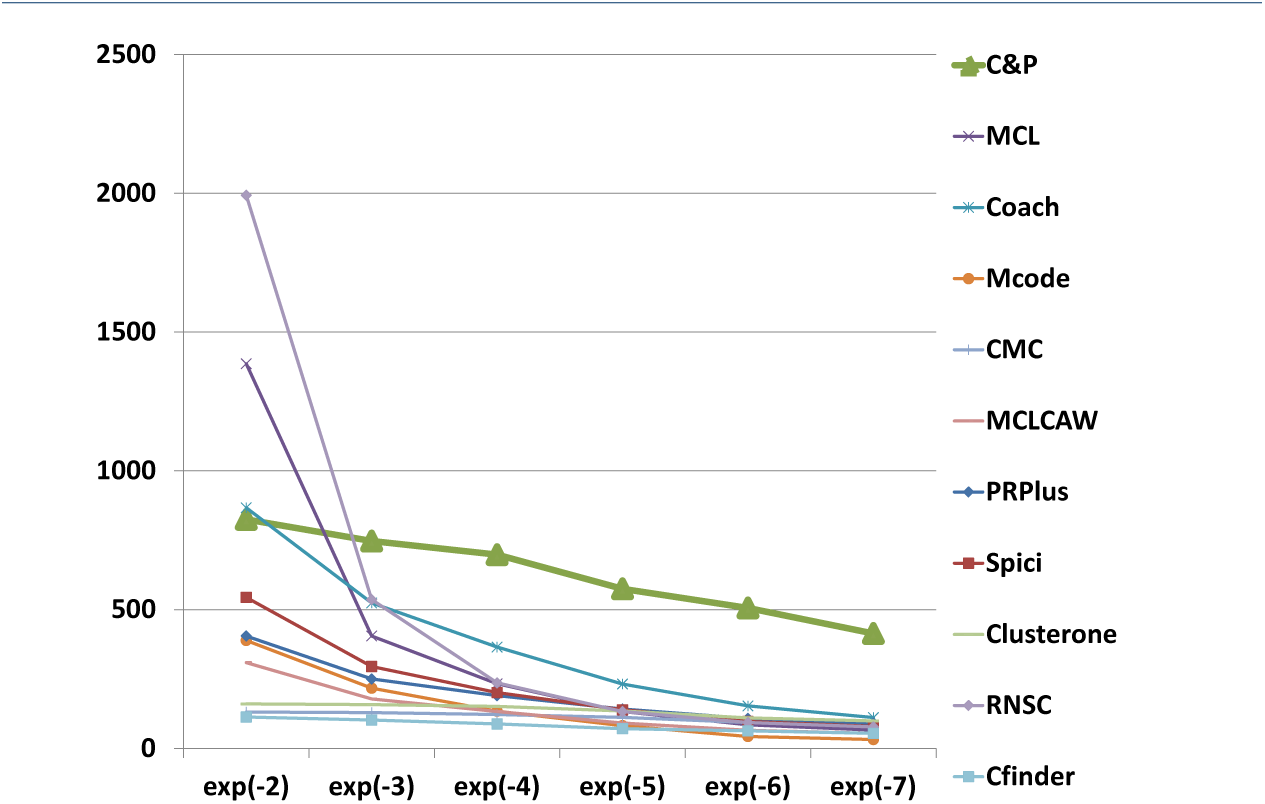
Number of predicted clusters with GO enrichment q-value below threshold, as a function of the threshold, for DIP yeast data.

**Figure 12.**
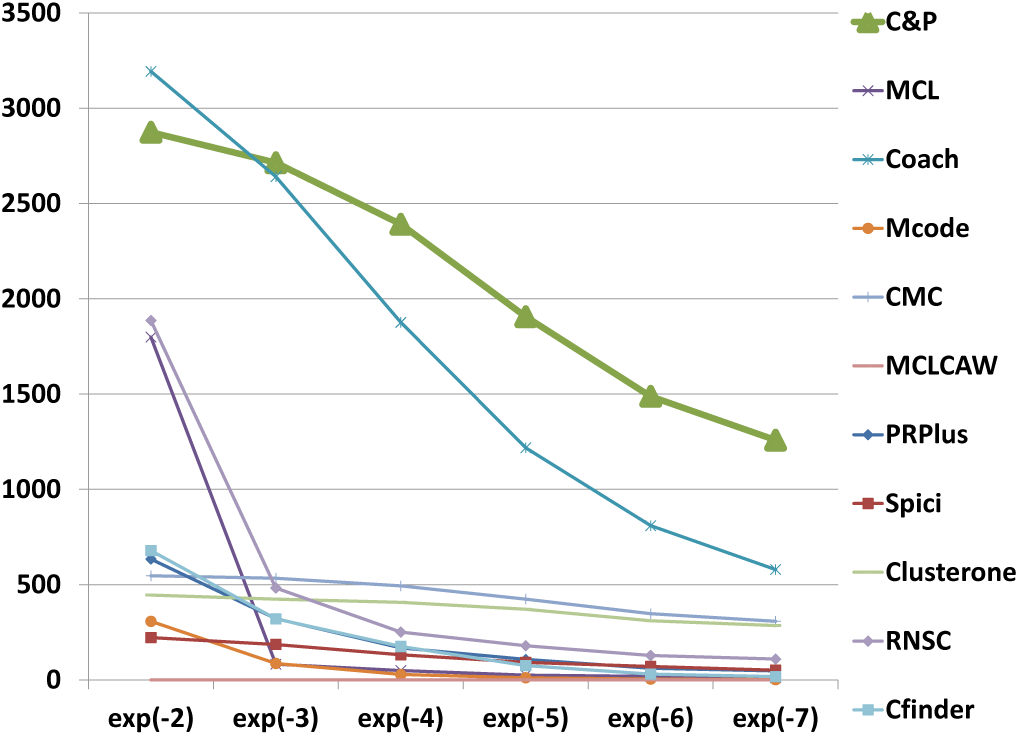
Number of predicted clusters with GO enrichment q-value below threshold, as a function of the threshold, for Biogrid yeast data.

**Figure 13.**
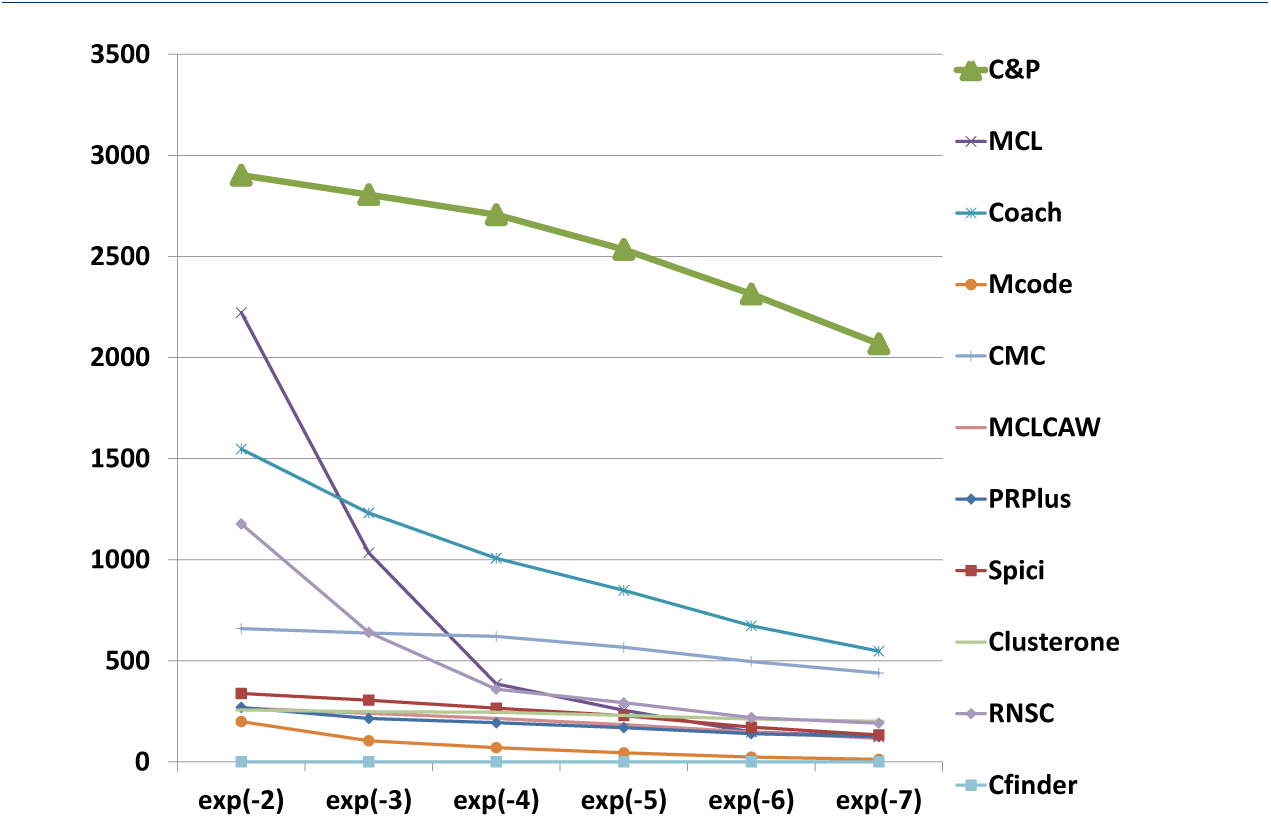
Number of predicted clusters with GO enrichment q-value below threshold, as a function of the threshold, for String yeast data.

**Figure 14.**
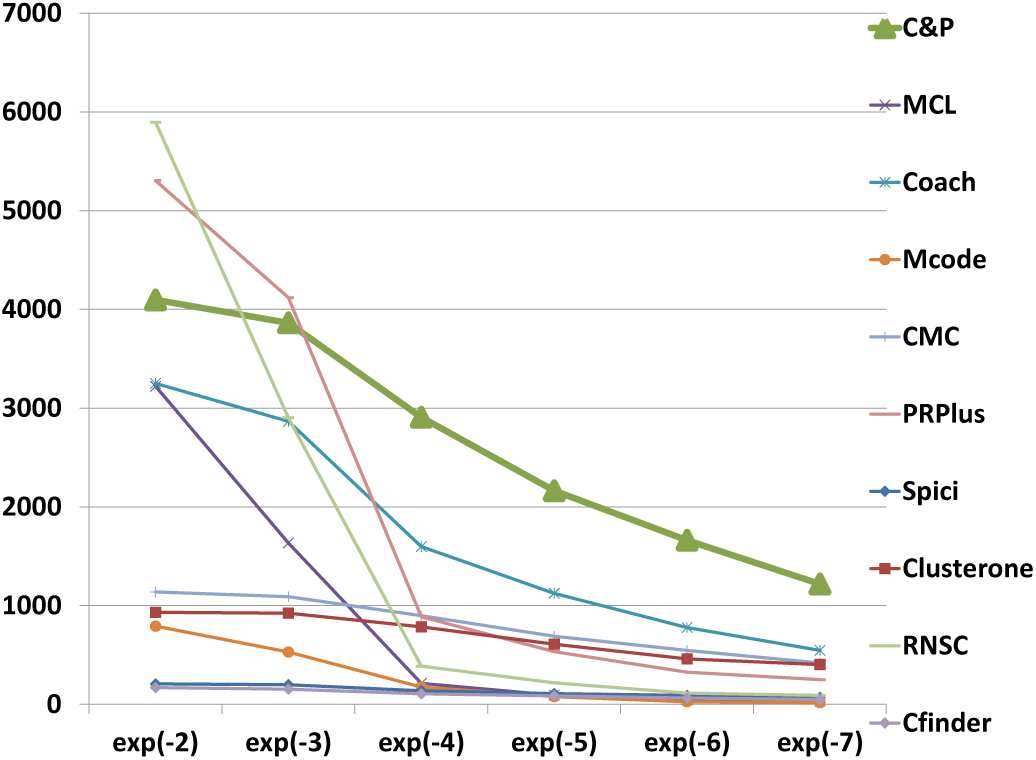
Number of predicted clusters with GO enrichment q-value below threshold, as a function of the threshold, for Biogrid Homo sapiens data.

**Figure 15.**
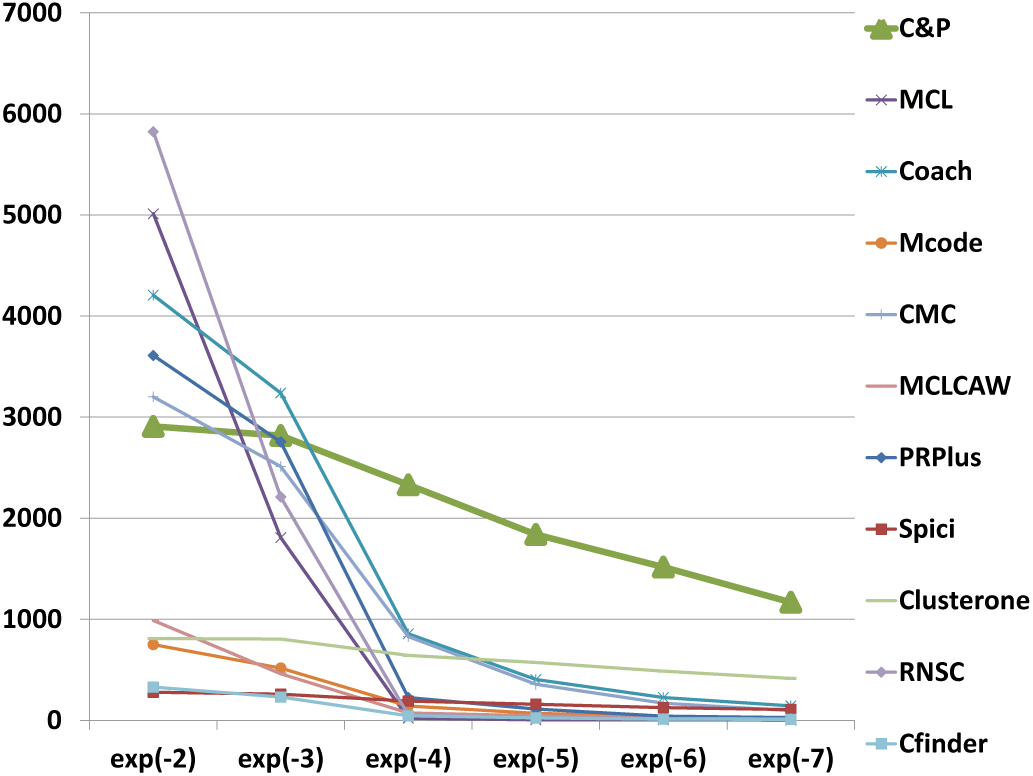
Number of predicted clusters with GO enrichment q-value below threshold, as a function of the threshold, for Biogrid Homo sapiens data (UBC removed).

**Figure 16.**
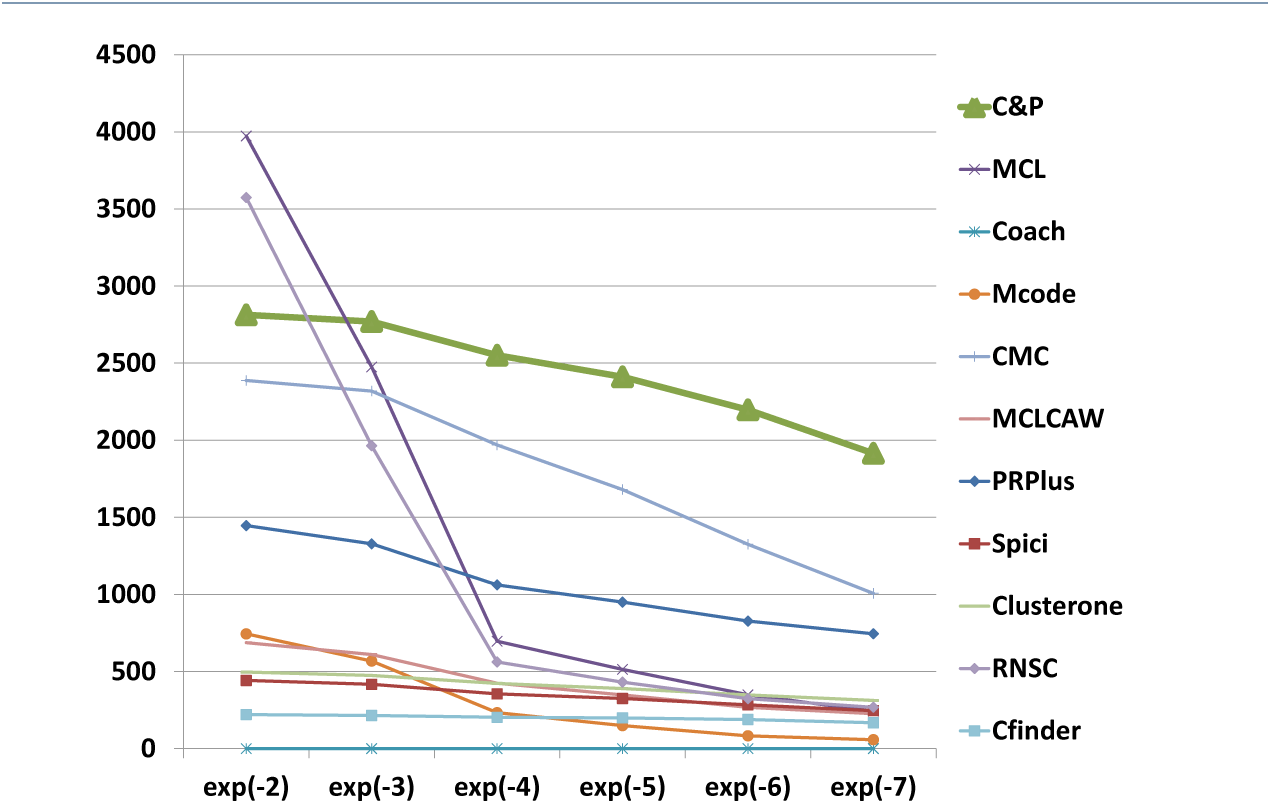
Number of predicted clusters with GO enrichment q-value below threshold, as a function of the threshold, for String Homo sapiens data.

*Core&Peel* has a larger or equal absolute number of high quality predictions below q-value 10^−3^ than the competing methods on five data sets out of six. For the BG-hs-UBC dataset *Core&Peel* leads below q-value 10^−4^. The overall trend is fairly consistent for all the six data sets tested.

Table 6 reports examples of predicted clusters with a notable low p-value, and the corresponding GO class. The top 10 predicted clusters we identify have *p*-value for their prevalent GO-annotation (all distinct) in the range 10^−191^ - 10^−72^. For a comparison, the top 10 functional modules detected by the recent method ADM (Adaptive Density Modularity) of Shen et al. [65, Table 3] relative to the same GO BP annotation have p-values in the range 10^−63^ - 10^−28^.

**Table 6.**
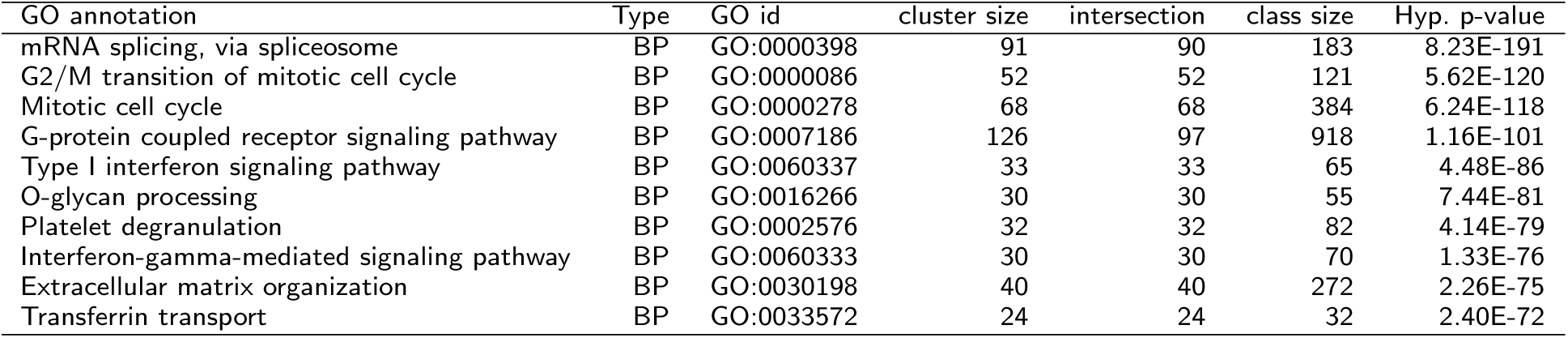
Data set STRING-CORE (homo sapiens) Functionally enriched clusters found with min size 8 and filtering policy 1. We report the top ten clusters by hypergeometric p-value. Each row reports: the GO annotation class, GO class type (BP=Biological Process), the GO id, the size of the cluster, the size of the intersection, the size of the functional class, and the hypergeometric p-value.

### 6.3 Robustness against noise in the PPIN graph

We have tested our method for its robustness against injection of random noise in the input network. Starting with the the Biogrid HS network we have added randomly additional (noise) edges for a number of additions ranging from 5% to 25% of the initial number of edges in steps of 5%. We have generated 10 networks for each class and taken the mean value of the 4 basic quality indices of section 4.2. The results are remarkably robust showing for three indices no variation up to the fourth decimal digit, and for the f-measure a variability of 0,001 across the range of noise values. Further tests with large random graphs are described in Additional_file_1 section 7, where we use the two stage multiple hypothesis test proposed in [66, 67] to bound the false discovery rate (FDR) associated with the identified complexes.

### 6.4 Running times

Figures 17, 18, 19, 20, 21, and 22 report in logarithmic scale the running times (seconds) for the 11 algorithms on the six data sets, with the parameters optimizing the f-measure. For MCL-caw we report the post-processing time in the graphic, thus a timing comparable with those of the other methods requires adding the MCL datum. Spici is the fastest method on all the data sets, often completing in less than a second. *Core&Peel* comes second in speed in all the data set (except for DIP where it is third).

**Figure 17.**
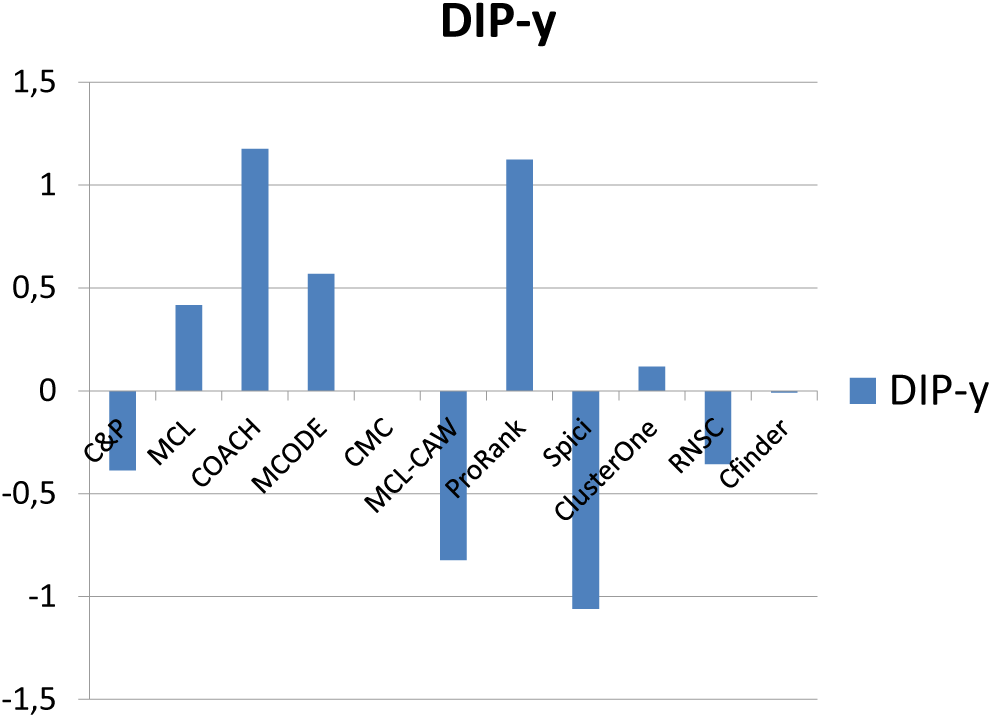
Time (in seconds) in Log10 scale for DIP data. Runs optimizing the f-measure for each algorithm.

**Figure 18.**
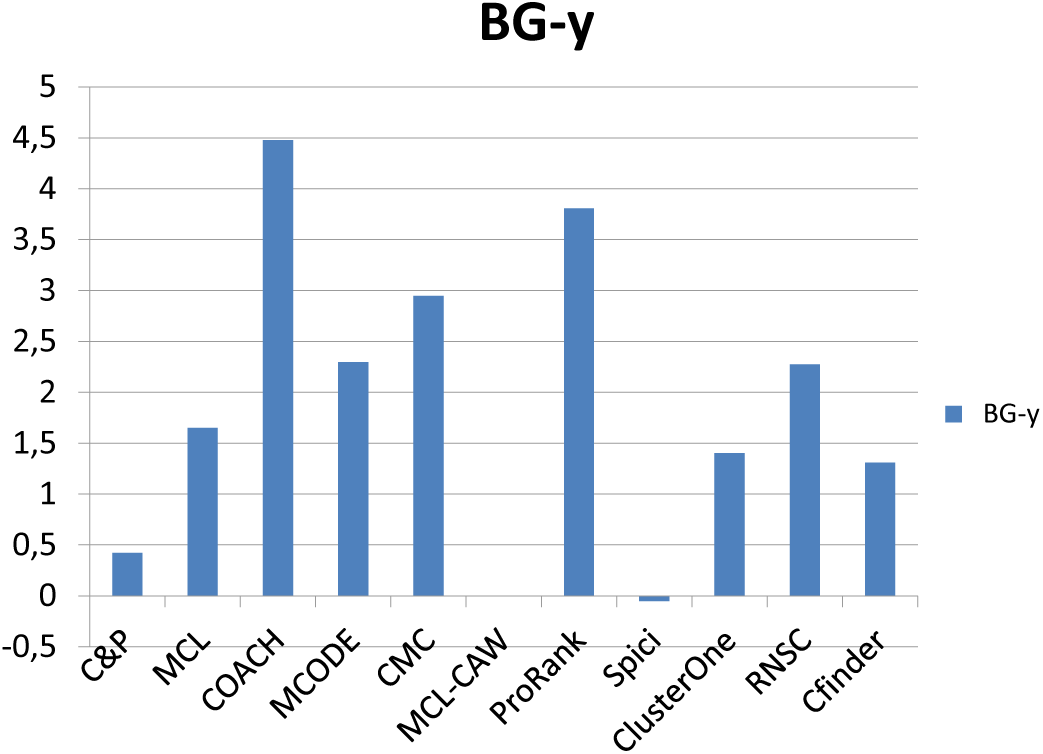
Time (in seconds) in Log10 scale for Biogrid yeast data. Runs optimizing the f-measure for each algorithm.

**Figure 19.**
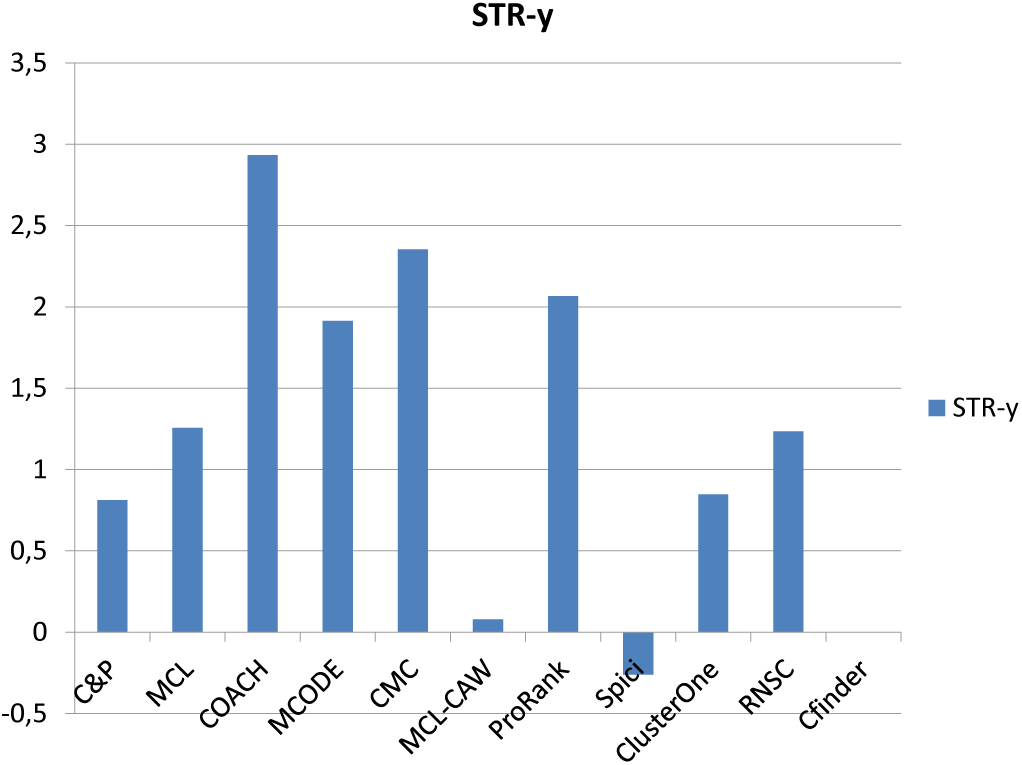
Time (in seconds) in Log10 scale for String yeast data. Runs optimizing the f-measure for each algorithm.

**Figure 20.**
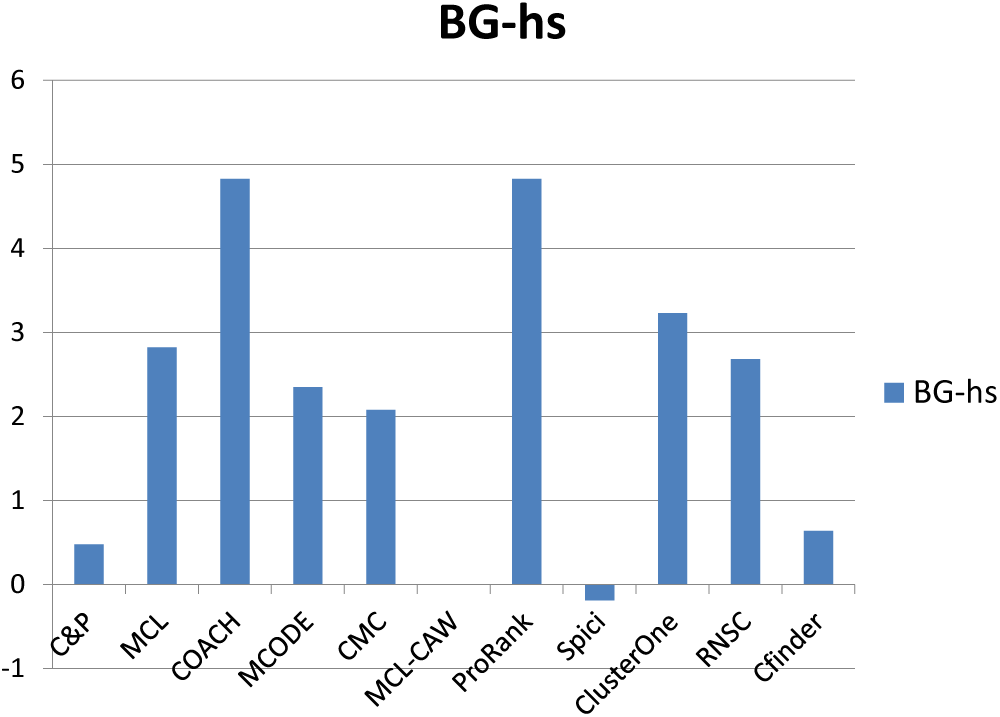
Time (in seconds) in Log10 scale for Biogrid homo sapiens data. Runs optimizing the f-measure for each algorithm.

**Figure 21.**
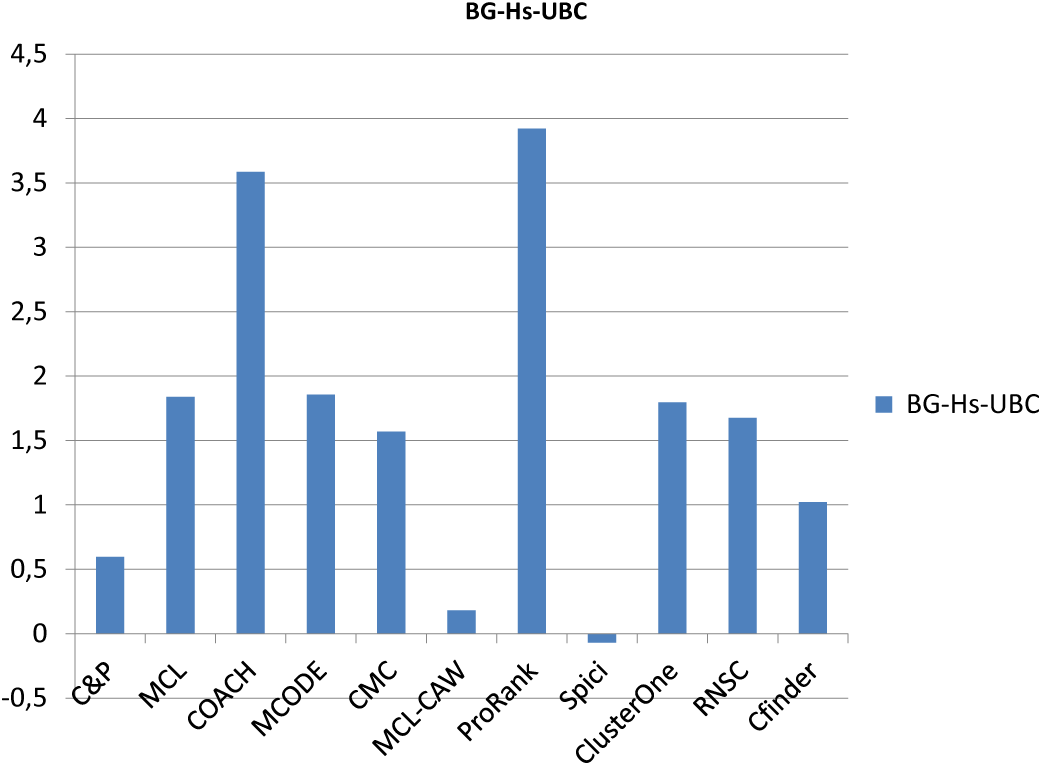
Time (in seconds) in Log10 scale for Biogrid homo sapiens data without UBC. Runs optimizing the f-measure for each algorithm.

**Figure 22.**
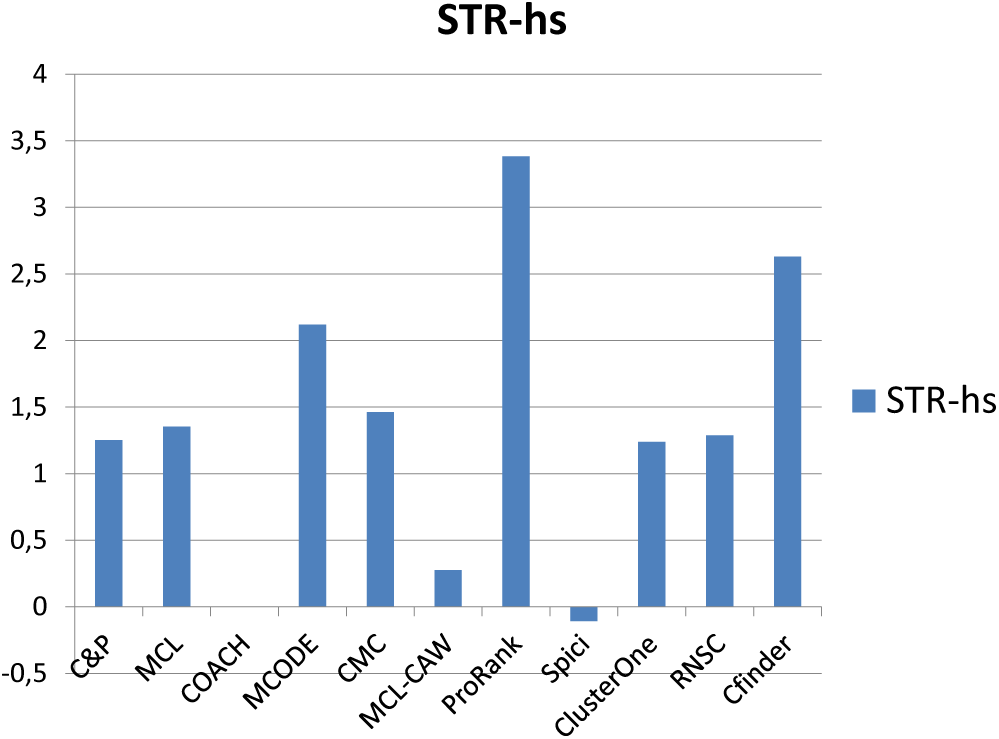
Time (in seconds) in Log10 scale for String homo sapiens data. Runs optimizing the f-measure for each algorithm.

### 6.5 Some predictions with support in the literature

In the long run the effectiveness of a protein-complex prediction method hinges upon its capability to uncover interesting and unexpected new phenomena of biological relevance. As an intermediate step we report on predictions made by *Core&Peel* that involve a known complex and one or two additional proteins, for which there is evidence of a biological function in the literature. We take the clusters detected by *Core&Peel* in the Biogrid and String homo sapiens network, we rank them by the highest value of the Jaccard correlation coefficient (jacc) and the semantic similarity (SS) with a matching known complex, we analyze the discrepancies (i.e. proteins non listed in the known complex but with a large number of PPI connections to it) and we highlight the literature supporting the functional relevance of the interaction. We chose here the parameter setting maximizing the f-measure.

Case 1 **(rank=2, jacc=0.875)**. This predicted cluster matches almost perfectly with the **20S proteasome** complex (Corum-id 191). Moreover the predicted cluster includes two additional proteins: **UBC** (Ubiquitin C) and **IQCB1** (IQ motif containing B1). The *hitpredict database* (http://hintdb.hgc.jp) also predicts high quality interactions between **IQCB1** and six **PSA** plus three **PSB** proteins. The functional connection of **UBC** and the **proteasome complex** within the protein degradation pathway is also well known (see e.g. [68]).
Case 2 **(rank=5, jacc =0.875)**. This predicted cluster matches almost perfectly with complex **TFIIH** (transcription factor complex TFIIH)(Corum-id 5495). Moreover the predicted cluster includes an additional protein: **AR** (androgen receptor). Indeed the phosphorilation action of **TFIIH** upon **AR** is reported in [69].
Case 3 **(rank=9, jacc=0.833)**. This predicted cluster matches almost perfectly with the **TFIIIC** containing complex (corum-id 1105), (and also with TFIIIC2, a second TFIIIC containing complex, corum-id 1101). Moreover the predicted cluster includes an additional protein: **GTF3C6** (general transcription factor IIIC, polypeptide 6). Dumay et al. [70] identified a sixth human **TFIIIC subunit**, specifically **GTF3C6**, which corresponds to a previously uncharacterized 213-amino acid human protein (C6ORF51).
Case 4 **(rank=14, jacc=0.823)**. This predicted cluster matches almost perfectly with the **20S proteasome** complex (Corum-id 191). Moreover the predicted cluster includes two additional proteins: PSMB8 (Proteasome subunit beta type-8) and **POMP** (proteasome maturation protein). The protein encoded by the **POMP** gene is a molecular chaperone that binds the **20S preproteasome** components and it is essential for **20S proteasome** formation. The POMP protein is degraded before the maturation of the **20S proteasome** is complete. A mutation in the 5′ UTR of this gene has been associated with KLICK syndrome, a rare skin disorder ([71]).
Case 5 **(rank 15, jacc = 0.818)**. This predicted cluster matches almost perfectly with PA700 complex (**26S protease/19S protease**) (corum-id 32). Moreover the predicted cluster includes an additional protein: **UCHL5** (ubiquitin carboxyl-terminal hydrolase L5). Interestingly, Darcy et al. [72] report that a small molecule (b-AP15) inhibits the activity of two **19S** deubiquitinases regulatory particles: ubiquitin C-terminal hydrolase 5 (**UCHL5**) and ubiquitin-specific peptidase 14 (USP14), resulting in accumulation of polyubiquitin, which in turn induces tumor cell apoptosis. Thus Darcy et al. suggest that the deubiquitinating activity of this regulatory molecule may form the basis for a new anticancer drug.
Case 6 **(rank=16, jacc=0.818)**. This predicted cluster matches almost perfectly with BAF complex (Synonyms: **SWI/SNF** complex A) (corum-id 1251 and 1237). Moreover the predicted cluster includes two additional proteins: **BCL7B** (B cell CLL/lymphoma 7B) and ARID1B (AT rich interactive domain 1B SWI1-like). A connection of the **SWI/SNF** complex with the first protein (**BCL7B**) is described in [73] where it is reported a proteomic analysis of endogenous **mSWI/SNF** complexes, which identified several new dedicated stable subunits of **SWI/SNF** complexes, including, among others, **BCL7B**.
Case 7 **(rank=20, jacc=0.8)**. This predicted cluster matches almost perfectly with corum-id 1097, **eIF3** complex (which is made of 13 proteins). Moreover the predicted cluster includes an additional protein: **Gag-Pol** [Human immunodeficiency virus 1]. It is reported in [74] that”A conserved structure within the HIV gag open reading frame that controls translation initiation directly recruits the 40S subunit and **eIF3**”.
Case 8 **(rank 27, ss=0.952)**. This predicted cluster matches almost perfectly with **SAP complex** (Sin3-associated protein complex) (corum id 591). Moreover the predicted cluster includes an additional protein: **ING2** (inhibitor of growth family, member 2). It is reported in [75] that “Besides the paralogous proteins, including HDAC1/HDAC2, mSin3A/mSin3B, and the histone-interacting RbAp46/RbAp48 proteins, the mammalian Rpd3L/Sin3L complex comprises at least five other sub-units, including **SAP30**, Sds3, SAP180/RBP1, SAP130, and ING1b/**ING2**, whose precise roles at the molecular level are poorly understood but most likely involve targeting the complex to specific genomic loci via one or more interaction surfaces”.
Case 9 **(rank 28, ss= 0.92)**. This predicted cluster matches almost perfectly with the **Ribosome** complex (corum-id 306). Moreover the predicted cluster includes an additional protein: SIRT7 (**Sirtuin 7**). Tsai et al. [76] investigate the role of **Sirtuin 7** in the **Ribosome** biogenesis and protein synthesis.
Case 10 **(String-hs data, rank=10, jacc=0.916)**. This predicted cluster matches almost perfectly with the **Exosome** (11 prot) complex (corum-id 789). Moreover the predicted cluster includes an additional protein: **XRN1** (5′-3′ exori-bonuclease 1). Li et al. [77] describe the competing role played by **XRN1** and the **Exosome** complex in Hepatitis C-Virus RNA decay.

## 7 Discussion

The experimental results reported in Section 5 show that *Core&Peel* is remarkably consistent in finding known complexes across 1 medium and 5 large data sets, ranking first in aggregated score against ten state-of-the-art methods in all 6 cases (CMC is second trice; SPICi twice, and Clusterone once).

*Core&Peel* also leads in the ability to produce cluster predictions that are highly consistent with GO-BP annotations. The specific complex-protein interaction predictions listed in section 6.5 have all a strong support in the literature. Although such predictions may not always correspond to actual complexes, they do indeed point at functionally relevant phenomena.

The *Core&Peel* algorithm exploits properties of complexes embedded in PPINs (egocentricity, density) that are more evident the larger the PPINs become, and it does not suffer from phenomena of combinatorial explosion (as both the theoretical analysis and the empirical running time attest). Thus we believe that *Core&Peel* can become a method of choice when even larger PPINs are built and analyzed, such as those arising in multi-species PPIN studies (see [37]) and those arising in immunology studies ( see [40]).

*Core&Peel* is fast and easy to use, requiring the setting of very few natural parameters relative to the minimum size, density and separation of the target complexes. Indeed, having a small sample of the type of complexes to be sought, these parameters can be extracted directly form the sample.

*Core&Peel* uses very little biological information except that embedded in the PPIN topology. Thus we believe further gains can be achieved by augmenting our scheme with the ability to handle PPINs endowed with edge weights^[12]^ (modeling, for example, PPI quality, or other types of a *priori* knowledge), or by incorporating GO annotations-based filters within the basic algorithmic framework. Improvements and tests along these lines are left for future research.

In this paper our main focus is to compare our proposed algorithm versus 10 competing algorithms on a sufficiently diverse pool of test data (2 species, 3 repositories, 6 PPIN, 2 PC golden standard sets) so to gain confidence in the robustness of the main thesis (i.e. the suitability of *Core&Peel* for discovering PC in large PPIN). We do not aim at suggesting that a particular type of PPIN repository should be preferred over others, and we do not even aim at implying that one should always use large PPIN in place of smaller ones (see e.g. [78]). Both questions are worthy of attention but fall outside the scope of the present article. The choice of the PPI data to be used for a given study is a non-trivial choice since many hidden biases could be implicit in the data (due both to its experimental origin, and to subsequent filtering) [79], thus these issues should be considered carefully at the initial stage of any experimental design.

## Author’s contributions

MP devised the algorithm and its analysis, drafted the manuscript, and exercised general supervision. MB collected the data, wrote the evaluation code and performed experiments. FG performed experiments and set up the web site interface. All authors read and approved the manuscript and contributed extensively to the work presented in this paper.

## Acknowledgments

The authors would like to thank the participants to the NII Shonan Meeting “Towards the ground truth Exact algorithms for bio-informatics research”, January 17-20, 2014. Work partially supported by the MIUR Flagship Project *Interomics*.

## Additional Files

Additional file 1 – Detailed Experimental settings

## Footnotes

[1] We drop *G* and *r* from the notation when they are clear from the context.

[2] BIOGRID-ORGANISM-Homo_sapiens-3.2.104.tab2.txt

[3] BIOGRID-ORGANISM-Saccharomyces_cerevisiae-3.2.104.tab2.txt

[4] protein.links.v9.05.txt.zip

[5] Scere20141001.txt

[6] Homo_sapines_gene_info.txt on 14/10/2013

[7] saccharomyces_cerevisiae_gene_info on 20/09/2013

[8] gene.association.goaJiuman.gz

[9] gene_association.sgd.gz

[10] Namely: MCL, Coach, MCODE, CMC, MCL-CAW, ProRank+, SPICi, Clus-terOne, RNSC, and Cfinder. A brief description of each is in Additional_file_1 section 1.

[11] Rolland et al. in [64] also remove interactions involving UBC in their high quality human PPIN.

[12] See discussion in Additional_file_1, Sec 5.

